# An Empirical Study on the Motivation of Helping Behavior in Rats

**DOI:** 10.1101/2023.02.01.526568

**Authors:** Shu Han, Ya-Qin Chen, Benhuiyuan Zheng, Ya-Xin Wang, Bin Yin

## Abstract

Using rodent models to explore the motivation of helping behaviors has become a new trend in recent years. Empathy, the alleviation of personal distress, and desire for social contact have been considered motivations for rodents to engage in helping behaviors. We used 108 Sprague-Dawley rats as subjects and modified the two-chambered helping behavior experimental setup in Carvalheiro and colleagues’ study to explore the main motivations of helping behavior in rodents through three experiments. The findings suggest that (1) the desire for social contact and pursuit of an interesting environment are the primary motivations for helping behavior, regardless of the presence of a dark chamber, and (2) the alleviation of personal distress and prior experience of social contact rather than distress experience contribute to the onset and persistence of helping behavior.

## 1 Introduction

Helping behavior, an important form of prosocial behavior, involves assisting others in solving their problems or alleviating their suffering (Stukas & Clary, 2012). It is oriented toward and beneficial for others (Mason, 2021). Helping behaviors are intentional by definition, and those behaviors that are unintentional but beneficial to others does not fit into this category (Mason, 2016). Helping behaviors can be instrumental or emotional (Bamberger et al., 2017). The former refers to providing specific tangible or goal-oriented help, such as donating and giving assistance, and the latter refers to providing emotional support, such as caring, comforting, and encouraging. Babbling infants, in the first and second years of life, try to understand the pain of others, and express concern through hugs and pats as gestures of emotional help (Zahn-Waxler et al., 1992). However, helping behavior is not unique to humans; nonhuman primates also express care and help for their kind. Bonobos comfort their companions with physical contact after seeing them experience a setback (Clay & de Waal, 2013a, 2013b). Lome, a chimpanzee, refused to enjoy a meal alone and shared it amicably with her companion (Schmelz et al., 2017). Two wild female bonobos actively adopted outgroup cubs (Tokuyama et al., 2021). Their helping behaviors transcend kinship.

It is not a surprise that nonhuman primates engage in helping behaviors. It seems implausible that relatively inferior mammals would do so. However, as early as in the 1960s, Rice and Gainer (1962) found that the free-ranging rat helped a trapped rat that was subjected to electric shock or suspension pressure in another chamber by pressing a lever. Free rats have tried to rescue their cagemates trapped in a restrainer (Bartal et al., 2011, 2014, 2016, Bartal 2021; Cox & Reichel, 2020; Ueno et al., 2019), or companions struggling in water chambers (Cox & Reichel, 2020; Kandis et al, 2018; Sato et al., 2015). They can distinguish between conspecifics and objects, as less door-opening rescue behavior manifests when they are confronted by toy rats and empty restrainers (Bartal et al., 2011). However, helping behavior continues to occur in the presence of “bystander rats” (Havlik et al., 2020). In the face of food temptations, they choose to avoid harming their companions and forgo “self-interest” (Hernandez-Lallement et al., 2020; Keysers et al., 2022). Rats prefer the side that provides food for both themselves and their companions to the side that provides food only for themselves (Márquez et al., 2015).

In these studies, helping behavior has been interpreted as the rodent’s ability to empathize with conspecifics in a state of distress, suggesting that free rats perceive the distress of conspecifics through an emotional contagion process and engage in helping behavior to alleviate such distress (Bartal et al., 2011, 2014; Cox & Reichel, 2020; Sato et al., 2015). Empathy is to the ability to perceive and experience the emotions of others and perform acts that benefit others (Huang & Su, 2012; Wang et al., 2021). However, Carvalheiro et al. (2019) argued that the motivation of the rodent’s helping behavior was to alleviate the pain they felt when they witnessed the suffering of their peers, and that the helping behavior would decrease when the free rat could escape the stressful situation. Lavery and Foley (1963) suspended a rat in the air to emit a painful scream, and found that the free rat spontaneously learned to press the crossbar either to reduce the pain of the suspended rat or to turn off the playback of the painful scream, thus terminating the aversive sound. Therefore, it was argued that the stressful state of the trapped rat can cause the free rat to feel distressed as well (Gonzalez-Liencres et al., 2014). When confronted with an event that causes aversion and pain, the rat resolves it by helping or escaping (Hernandez-Lallement et al., 2020; Knapska et al, 2010). There is also a view that rodents, as pack animals, prefer to interact with their own kind (Bibb et al., 1972) and that social contact and interaction are motivations for helping behavior (Hachiga et al., 2018; Hiura et al., 2018). Silberberg et al. (2014) suggested that when free and trapped rats could not interact socially, the waiting time for free rats to open the door became longer as the number of tests increased. This suggests that the inability to socially engage may hinder the occurrence of helping behaviors (Heslin & Brown, 2021; Schwartz et al., 2017). Views on the causes of helping behavior in rodents include both altruistic-motivated explanations of empathy and individualistic-motivated explanations of alleviating self-distress or pursuing social contact. However, there are still disputes over whether rodents are motivated by one of these factors or a combination of them, and to what extent the different factors play a role. this study tested the plausibility of these explanations by improving the experimental setup in Carvalheiro et al. (2019) and controlling whether rats could socially engage and escape a situation to alleviate their own distress.

Helper-recipient familiarity may influence helping behavior in rats. The fear of trapped rats can be transmitted to free rats only if the former are familiar with the latter (Gonzalez-Liencres et al., 2014). Free rats are quicker to help familiar than unfamiliar rats when faced with trapped rats with different levels of familiarity (Bartal et al., 2014). Burkett et al. (2016) supports this result. The current study, referring to Bartal et al. (2011, 2014), examined the phenomenon by manipulating different restrainer conditions (empty restrainers, toys rats, familiar and unfamiliar rats). Studies have found that experience being trapped can promote helping behavior in rats (Hernandez-Lallement et al., 2020; Sato et al., 2015), and experience of social contact can influence helping behavioral decisions in rats (Bartal et al., 2014). However, how these factors, in situations where escape from help is possible, influence helping behavior has not been explored. We experimentally explore this by applying dark chamber situations and reversing roles for free and trapped rats.

This study tested whether the following hypotheses are true: (1) Empathy is a motivation for helping behavior in rats and free rats have a shorter latency in opening the door when they are in a restrainer with their conspecific (including familiar and unfamiliar rats) than when they are in a restrainer with an object (including empty restrainers and toy rats). (2) Social contact is a motivation for helping behavior, and the door-opening latency is shorter when there is social contact with trapped rats. (3) Alleviating personal distress is motivation for helping behavior, and the door-opening latency is shorter when there is a dark chamber to alleviate personal distress. (4) The door-opening latency is shorter for free rats with previous trapping and social contact experience. (5) Familiarity affects the helping behavior of rats, and the door-opening latency is shorter when there are familiar trapped rats in the restrainer as opposed to when there are unfamiliar trapped rats. In sum, this study explores the main motivations for helping behavior through a comparative study of rodents and helps expand the scientific theoretical explanation of helping behavior motivation.

## 2 Methods

### 2.1 Subjects

A total of 108 4-month-old male Sprague-Dawley rats were used as subjects (54 each in Experiments 1 and 2, and subjects from Experiments 1 and 2 were used in Experiment 3). In Experiment 1, all subjects were randomly assigned to 18 rearing cages, with 3 per cage. Subjects were randomly divided into the experimental group (i.e., social contact condition, *n =* 21, containing 7 familiar trapped rats), control group (i.e., non-social contact condition, *n* = 21, containing 7 familiar trapped rats), and randomly determined as free (*n* = 2) and familiar trapped (*n* = 1) rats for each cage according to whether free rats could have social contact with trapped rats after opening the door during the experiment. Unfamiliar trapped rats (*n* = 12) constituted another subject in the experiment. In Experiment 2, the subjects were assigned in exactly the same way as in Experiment 1. In Experiment 3, the roles were reversed and the trapped rats (*n =* 28) from Experiments 1 and 2 were used as free rats, of which 14 each had and did not have social contact experience. Further, 14 from each group were divided into social and non-social contact conditions, according to whether they could have social contact with the trapped rats in Experiment 3, in order to obtain the social contact experience - social contact condition (*n* = 7), social contact experience - non-social contact condition (*n* = 7), non-social contact experience - social contact condition (*n* = 7), and non-social contact experience - non-social contact condition (*n* = 7). Ambient temperature (22 ± 1)°C, humidity (50 ± 5%), and light hours –21:00 - 9:00 the following day) were controlled, and food (20g/per rat) and water were administered regularly on a daily basis.

### 2.2 Tools and materials

To investigate how factors such as desire for social contact, alleviation of self-distress, and previous trapping experience affect helping behavior in rats, the present study modified the experimental setup in Carvalheiro et al. (2019) by adding a middle chamber to control whether free rats could socially engage with trapped rats. As seen in **Fig. 1**, from left to right, the three chambers (each 40 × 40 × 60 cm) are the trapped, middle, and dark ones. The restrainer (20 × 10 × 5 cm) was placed in the trapped chamber, and after the trapped rat was placed inside, the free rat could discontinuously touch the touch sensor (FR - 5 discontinuity) on the wall of the middle chamber, and the door of the restrainer opened and the trapped rat was rescued. The transparent walls of the trapped room and the middle chamber allowed visual information exchange, the area where the walls of the restrainer and middle chamber were connected had sniff holes to allow auditory and olfactory information transfer, and the middle chamber was separated from the dark chamber by opaque black walls. Whether door 1 opened or not determined whether the free rats could have social contact with the trapped rats and was controlled by the computer program Graphic State 4 (Coulbourn Instruments, Holliston, MA) based on the experimental conditions. Whether door 2 opened or not determined whether the free rats could enter the dark chamber to escape from the help situation and was controlled manually based on the experimental conditions.

**Figure 1.**
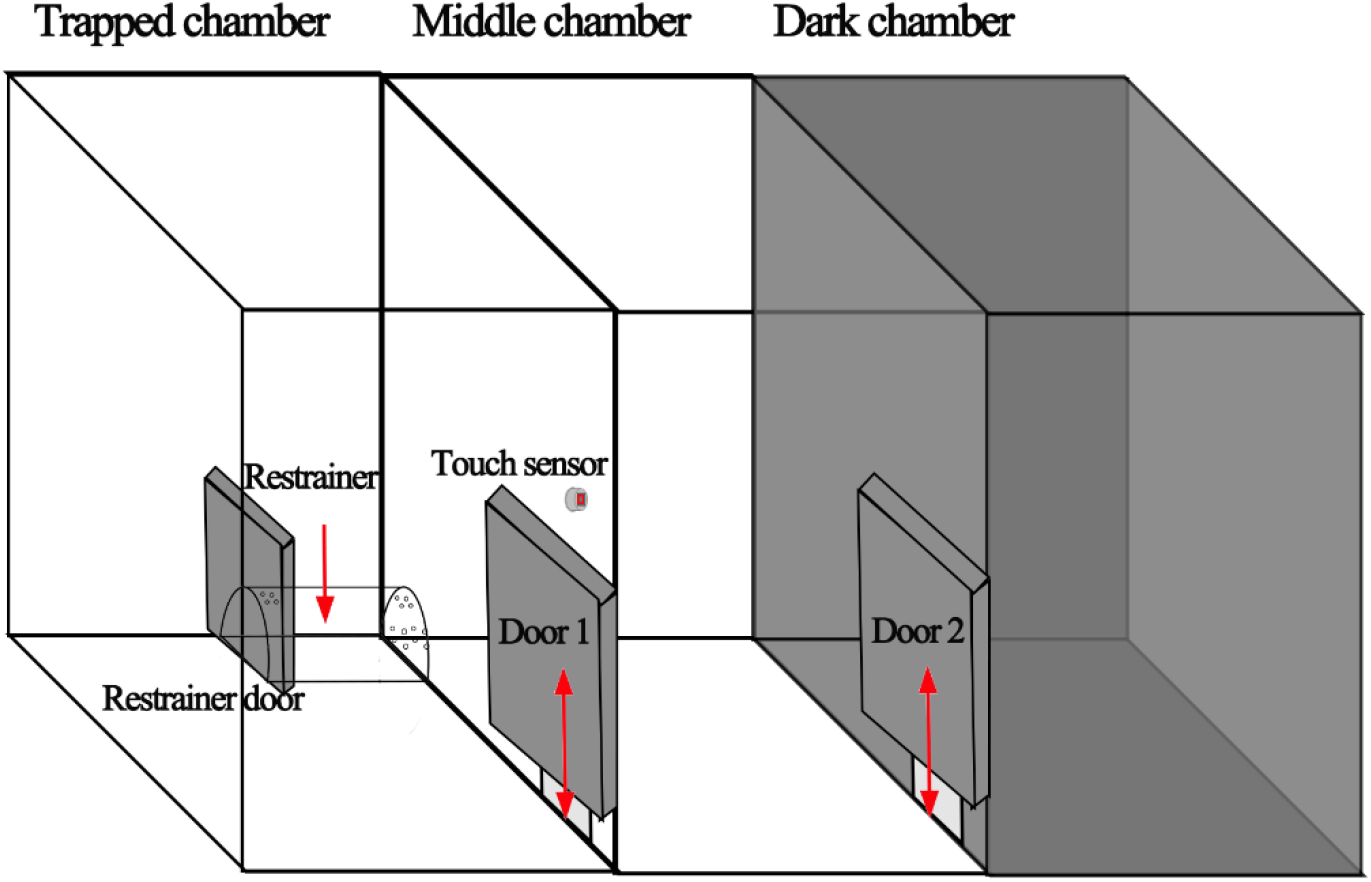
Experimental setup in this study

### 2.3 Experimental design

In this study, the restrainer door-opening paradigm was used to put trapped rats subjected to restrainer stress in a state of discomfort (Campos et al., 2010; Pitman et al., 1988) to explore the door-opening helping behavior of free rats. There were three experimental designs. In Experiment 1, a two-factor mixed design of 2 (social contact: allowed, not allowed) × 4 (restrainer condition: empty restrainer, familiar rat, unfamiliar rat, toy rat) was used to investigate the effect of social contact on helping behavior of rats in an inability-to-escape-context, using the ability to have social contact with trapped rats and the restrainer condition as between- and within-subjects variables, respectively. The design of Experiment 2 was the same as that of Experiment 1, but provided a place for subjects to escape from the helping situation (dark chamber). Experiment 2 investigated the effect of relieving one’s pain by escaping the situation on the helping behavior of rats. Experiment 3 reversed the roles of free and trapped rats in Experiments 1 and 2, and conducted 2 (previous social contact experience when trapped: yes, no) × 2 (social contact: allowed, not allowed) × 4 (restrainer conditions: empty restrainer, and familiar, unfamiliar, toy rat) experiments under the condition that both free rats had trapping experience in a three-factor mixed design, with prior social contact experience at the time of trapping and whether or not social contact with the trapped rat was allowed during the current experiment as between-subject variables and the restrainer condition as a within-subject variable, to investigate the effects of prior trapping or social contact experience on the helping behavior of rats through role reversal. Four restrainer conditions were set in all three experiments to explore the effect of familiarity on helping behavior in rats.

### 2.4 Experimental procedures

1. Adaptation to the environment All rats were subjected to touch interaction for 5 minutes each for 11 days after they were enrolled in the storeroom to help them reduce their anxiety levels (Costa et al., 2012), adapt to the housing environment, get used to human contact, and establish good relationships with the main subject (as rats from Experiments 1 and 2 were used in Experiment 3, there was no need to re-touch the subjects in Experiment 3).
2. Adaptation to the experimental setup To reduce tensions among the rats during the test owing to the novelty of the environment, adaptation to the experimental setup was performed after adaptation to the environment. Door 1 and the restrainer door were opened to allow free rats to explore the device for 15 minutes. Adaptation of trapped rats took place before the testing phase began.
3. Door-opening learning phase After 24 hours, the door-opening learning phase of the free rats began. The feeding amount of rats was reduced on a daily basis, chocolate pills were placed inside the restrainer, and the semi-starved free rats were placed in the center of the middle chamber. They could explore it in such a way that if they touched the sensor on the wall, the door of the restrainer and door 1 would open simultaneously and they could enter the restrainer to enjoy the chocolate pills. To help the free rats that opened the door accidentally learn to do so purposefully, the door-opening learning was divided into three stages with increasing difficulty. In Stage 1, the free rats only needed to touch the sensor once (FR-1) for door 1 and the restrainer door to open. In Stage 2, the free rats needed to touch the sensor five times in a row (FR-5 continuously) for door 1 and the restrainer door to open. In Stage 3, the free rats needed to touch the sensor five times discontinuously (FR-5 discontinuously) for door 1 and the restrainer door to open. The operational definitions related to free rat behavior in the three stages are presented in **Table 1**. Learning ended when all free rat were able to maintain a low door-opening latency (≤ 200 s) for three consecutive trials in stage 3 (in Experiment 3, owing to experimental schedule constraints, learning ended after only one day of meeting the condition, but it was possible to ensure that the free rats had learned to open the door). Those free rats who did not learn to open the door were given extra practice (chocolate was applied to the sensor to increase the probability of touching it), and those that consistently failed to learn to open the door were removed from the study. All subjects in this study successfully learned to open the door.

**Table 1.** Operational definitions of different conditions in the door-opening learning phase on rat behavior
4. Helpful behavior testing phase

| Stage | Conditions | Operational Definition |
| --- | --- | --- |
| **Stage 1** | Touching the sensor once | Any part of the body such as the head, nose, and front paw of the free rat touches the sensor once. |
| **Stage 2** | Touching the sensor five times continuously | The free rat's head, nose, front paws and other body parts touch the sensor, five times in a row, and there is no time interval required between each time. |
| **Stage 3** | Touching the sensor five inconsecutive times in a row | The free rat's head, nose, front paws and other body parts touches the sensor, five inconsecutive times in a row, with the time interval between each touching exceeding $\geq 1$ s. |

After ensuring that all free rats had learned to open the door and all trapped rats had adapted to the device, the helping behavior test began. The door-opening latency time(s) of the free rats was recorded, that is, the point from the time they entered the middle chamber to the time they successfully opened the door by not touching the sensor five times consecutively. It was found that the optimal upper limit of the door-opening time for each trial was 15 minutes, and if the door was not opened within 15 minutes, the door-opening latency was recorded as 900 seconds. In the social contact group, the free rats could enter the trapped chamber and explore and interact with the trapped rat after opening the door. In the non-social contact group, the free rats could only move around in the middle (Experiment 1) and dark (Experiments 2 and 3) chambers after opening the door.

This study was reviewed and approved by the Laboratory Animal Ethics Review Committee of Fujian Normal University (No.: IACUC-20210040) and were in accordance with the *National Research Council’s Guide for the Care and Use of Laboratory Animals*.

### 2.5 Experimental data processing

*Jamovi* was used to conduct multivariate repeated measure variance analysis and independent sample t test on the experimental results. Descriptive statistics were conducted on the door-opening latency (learning phase, helping behavior test phase, etc.) of the social and non-social contact groups under each condition, the significant differences among the conditions were tested (*p* < 0.05 was designated as statistically significant), and multiple comparisons and simple effect analysis were conducted. Graphpad Prism 8.0 was used for plotting. This dataset has been uploaded on the Science Databank of China and can be accessed at https://doi.org/10.57760/sciencedb.o00115.00014 for review.

## 3 Results

### 3.1 Learning Phase Results

Repeated ANOVAs conducted with experimental treatment and number of studies were the between- and within-group variables, respectively. The results of Experiment 1 (**Fig. 2, Table S1**) indicated that the between-group main effect was not significant *F*(1, 26) = 0.36, *p* = .553, *η*^2^ _*partial*_ = .01, thus showing that the difference in door-opening latency between the social and non-social contact groups in the learning phase was not significant. The results of Experiment 2 (**Fig. 3, Table S2-1)** showed a significant main effect between groups, *F*(1, 26) = 6.02, *p =* .021, *η*^2^ _*partial*_ = .19, and upon comparison, the difference was caused by four rats in the non-social contact group not opening the door on the first day of learning, but in the subsequent (except for the first day) learning (**Table S2-2**), the between-group effect was not significant, *F*(1, 26) = 3.38, *p* = .078, *η*^2^ _*partial*_ = .12, indicating that the difference in door-opening latency between the social and non-social contact groups in the learning phase was not significant.

**Figure 2.**
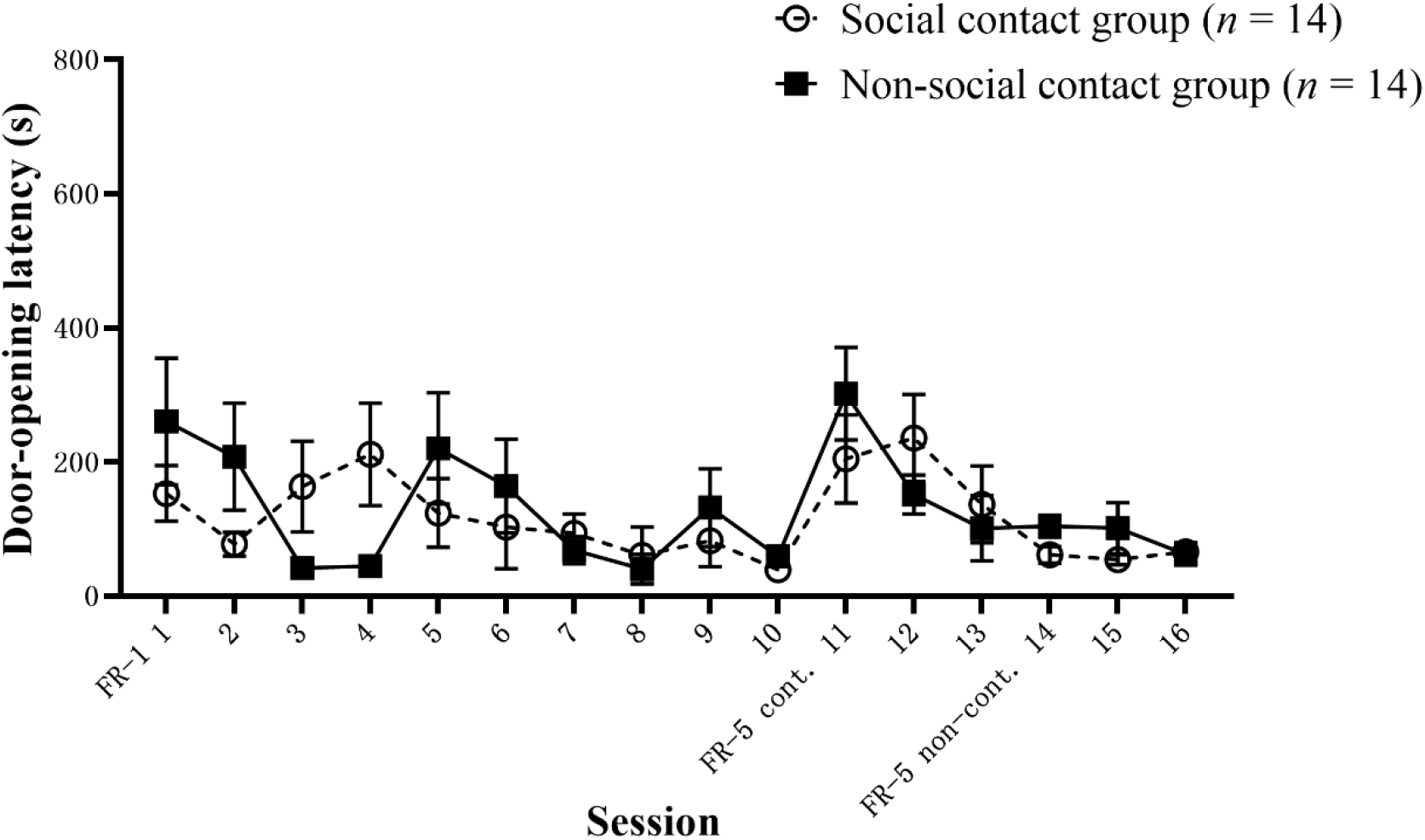
Door-opening latency in the learning phase in Experiment 1

**Figure 3.**
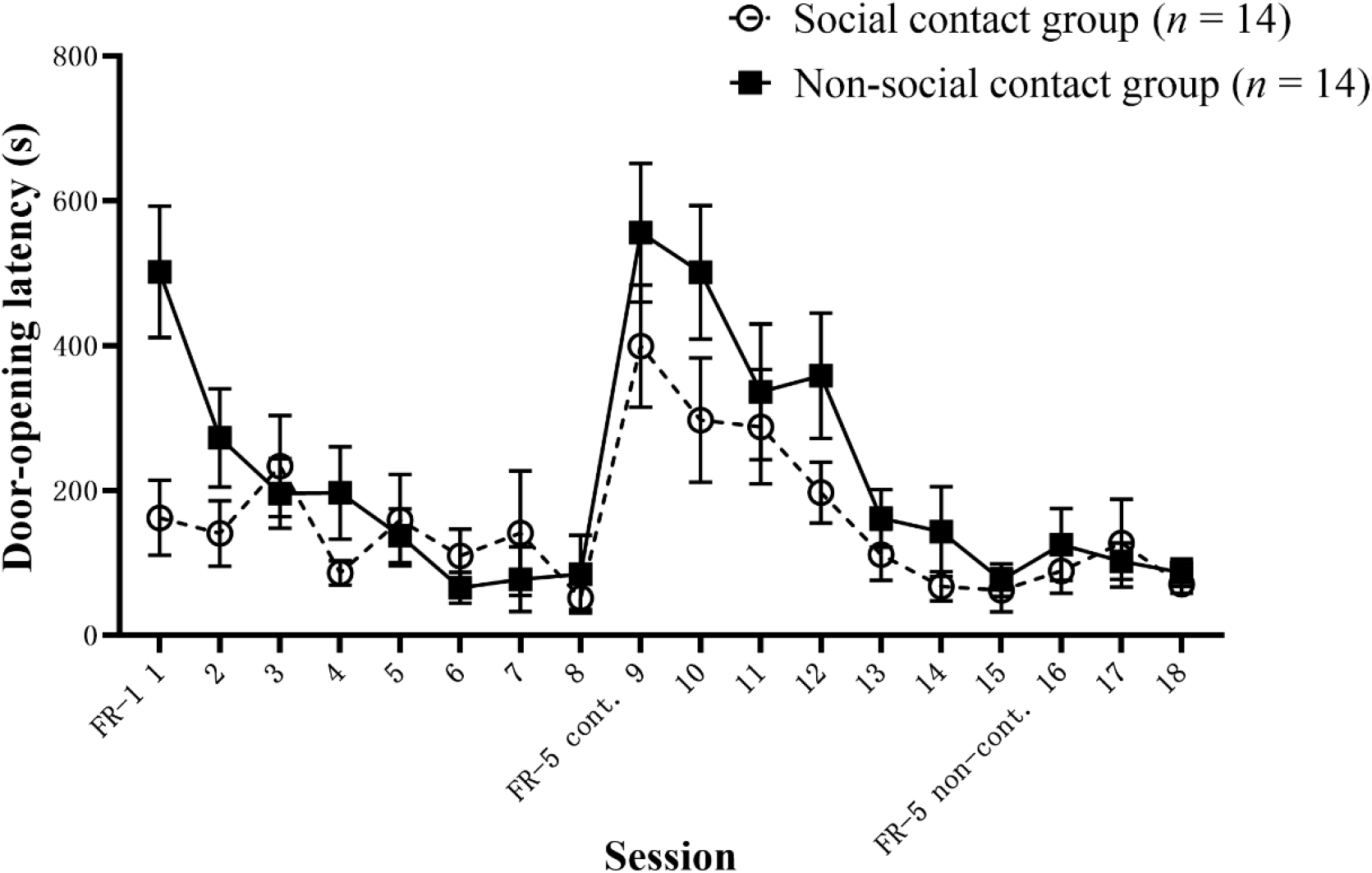
Door-opening latency in the learning phase in Experiment 2

In Experiment 3, repeated ANOVAs were conducted with the presence or absence of prior social contact experience and availability of social contact during the experiment, and the number of studies as between- and within-group variables, respectively. The results (**Fig. 4, Table S3**) showed that the main effect of prior social contact experience was not significant, *F*(1, 24) = 0.19, *p* = .669, *η*^2^ _*partial*_ = .01. The main effect of whether social contact was possible during the experiment was not significant, *F*(1, 24) = 0.52, *p* = .476, *η*^2^ _*partial*_ = .02, indicating that subjects in different experimental treatments had a learning phase. The difference in door-opening latency was not significant. In each experiment, not only did all free rats learn to open the door, but rats with different experimental treatments learned to open the door to an almost equal degree.

**Figure 4.**
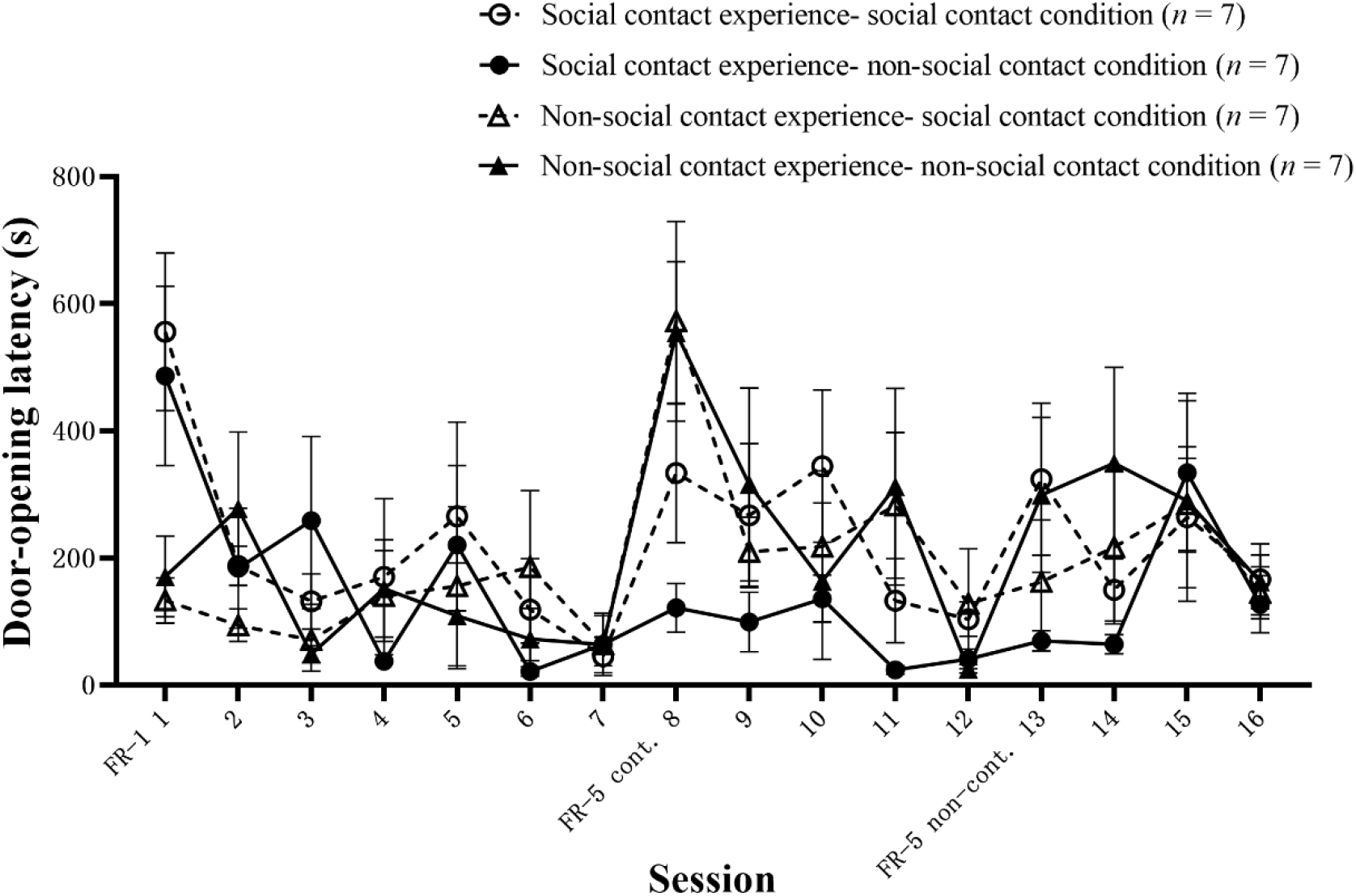
Door-opening latency in the learning phase in Experiment 3

### 3.2 Test phase results

#### 3.2.1 Experiment 1: The effect of social contact on helping behavior in the inability to escape condition

The helping behavior test in Experiment 1 was conducted over nine days, with each three-day period averaged and divided into early-, middle-, and late-periods in the order of the number of days tested (**Fig. 5**). The results of the ANOVA on door-opening latency (**Table S4-S5**) showed that the main effect of the experimental treatment was significant, *F*(1, 26) = 144.16, *p* < .001, *η*^2^ _*partial*_ = .85, and the door-opening latency was shorter in the social contact group (61.7±5.9 s) than in the non-social contact group (522.8±38.0 s), which supports hypothesis (2) and indicates that social contact is an important motivation for maintaining helping behavior in rats, and that free rats have shorter door-opening latencies when they can socially engage with trapped rats. The main effect in the test period was significant, *F*(2, 52) = 60.89, *p <* .001, *η*^2^ _*partial*_ = .70, and the door-opening latency was significantly shorter in the early-test period (175.5±28.8 s) than in the middle- (293.6±53.3 s) and late-test (407.7±69.1 s) periods, whereas the interaction between the test period and experimental treatment was also significant, *F*(2, 52) = 64.16, *p* < .001, *η*^2^ _*partial*_ = .71. There was no significant difference within the social contact group under different test periods (67.4±10.5 s, 56.1±5,9 s and 61.6±6.9 s for the early-, middle- and late-test periods, respectively), whereas the non-social contact group had a significantly shorter door-opening latency in the early-test (283.6±39.3 s) than in the middle- (531.0±55.8 s) and late-test (753.8±36.7 s) periods, which again support hypothesis (2) and indicate that when social contact is unavailable, the door-opening latency of the non-social contact group will gradually increase with testing experiences. The main effect of the restrainer condition was significant, *F*(3, 78) = 3.22, *p =* .027, *η*^2^ _*partial*_ = .11, but the door-opening latency in the object condition (278.8±46.2 s) was shorter than that in the similar condition (305.7±50.8 s), which does not support hypothesis (1). The difference in the door-opening latency between the familiar and unfamiliar rat conditions was not significant, *t*(26)= 0.38, *p* = 0.980, thus hypothesis (5) was not supported.

**Figure 5.**
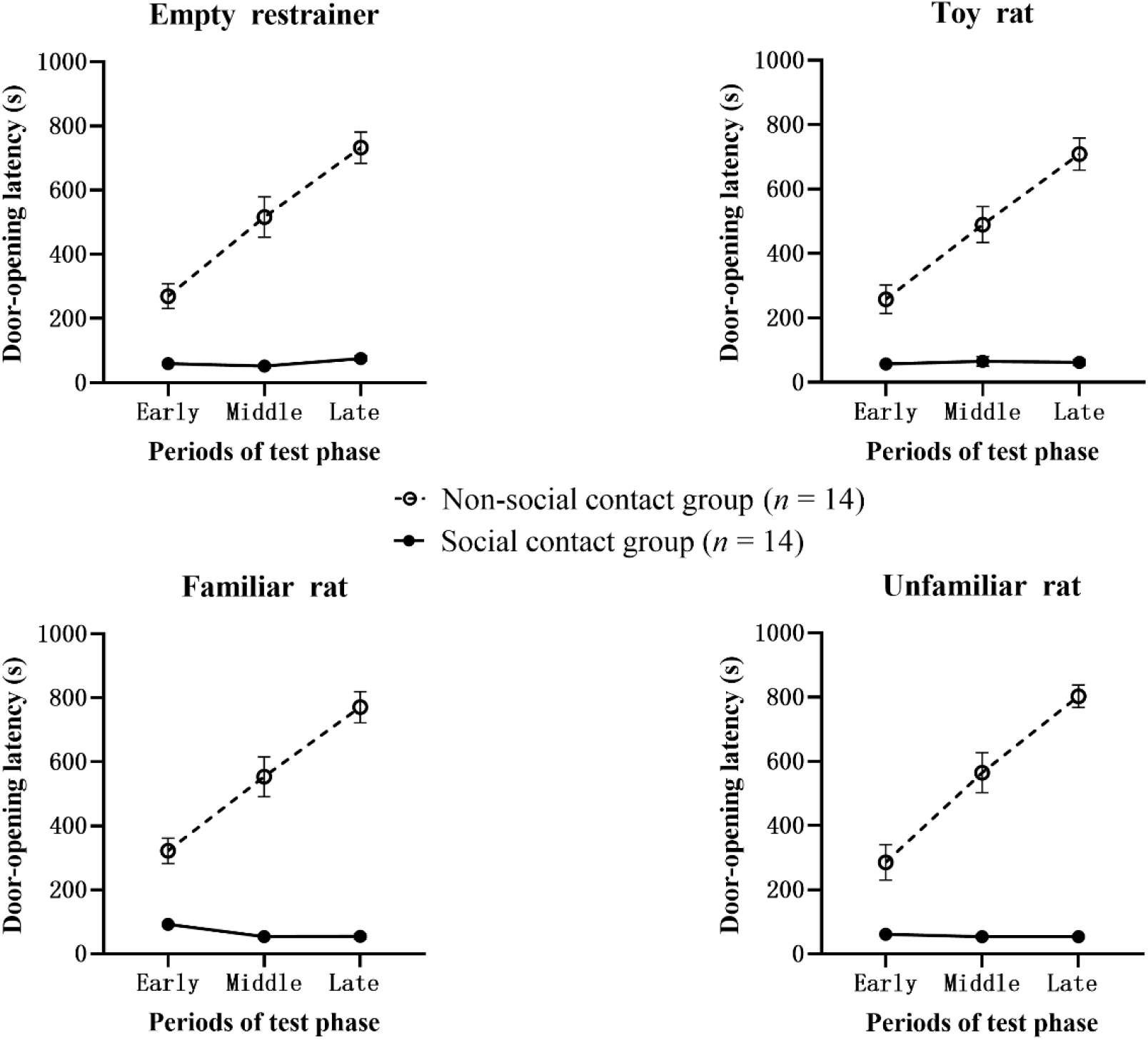
Door-opening latency in the test phase in Experiment 1

To further test the empathic properties of door-opening behavior among free rats, three cases from the social contact group in Experiment 1 were randomly selected according to qualitative coding (and two additional cases were added to determine theoretical saturation). The action sequences of free rats before opening the door in 180 videos were manually observed and coded, and it was found that there were similarities in the movement trajectories of different individuals before opening the door under the same restrainer condition (**Table 2**).

**Table 2.** The trajectory of the free rats before opening the door in Experiment 1

| Test phase | Empty restrainer | Familiar trapped rats | Unfamiliar trapped rats | Toy trapped rats |
| --- | --- | --- | --- | --- |
| **Early phase** | Walk-single touch-sniff-walk-single touch | Sniff-walk-single touch | Sniff-walk-sniffing-walk-single touch | Walk-single touch-sniff-walk-single touch |
|  | Walk-single touch-sniff-continuous touch | Sniff-walk-sniff-walk-single touch | Sniff-walk-single touch | Walk-single touch-continuous touch |
|  | Sniff-walk-single touch - continuous touch | Sniff-single touch-sniff-walk-single touch | Walk-sniff-sniff-walk-single-touch | Sniff-walk-single touch |
| **Middle phase** | Walk-single touch-sniff-walk-single touch | Sniff-walk-single touch | Sniff-walk-sniff-walk-single touch | Sniff-walk-single touch |
|  | Walk-sniff-single touch | Sniff-walk-single touch-continuous touch | Sniff-walk-single touch-continuous touch | Walk-single touch-sniff-walk-single touch |
|  | Walk-single touch-walk | Sniff-single touch-walk-continuous touch | Walk-sniff-walk-single touch | Walk-single touch - continuous touch |
| **Late phase** | Walk-single touch-sniff-walk-single touch | Sniff-walk-single touch | Sniff-walk-sniff-walk-single touch | Walk-single touch-sniff-walk-single touch |
|  | Walk-sniff-single touch | Sniff-walk-sniff-walk-single touch | Sniff-walk-sniff-walk - continuous touch | Sniff-walk-single touch |
|  | Sniff-walk-single touch | Walk-sniff-single touch-sniff-continuous touch | Walk-sniff-walk-single touch | Walk-sniff-single touch |

Notes: Sniff: sniff the hole; Walk: walk around the perimeter/center of the center chamber; Single touch: touch once at the sensor; Continuous touch: touch multiple times in a row at the sensor.

First, in the empty reatrainer condition, the free rats usually performed the movement sequence of “walk-single touch-sniff-walk-single touch,” that is, in each trial, the rats always walked around the middle chamber first after entering the device, instead of sniffing the holes first, and the movement trajectory in the toy rat condition was similar to that in the empty restrainer. In contrast, in the familiar and unfamiliar rat conditions, free rats usually performed the action sequence of “sniff-walk-single touch” or “sniff-walk-sniff-walk-single touch,” that is, free rats always sniffed the sniff hole first after entering the device, which may reflect their concerns for their trapped counterparts. In contrast, under the empty restrainer condition, the “continuously touch sensor” action appeared more and earlier in the early stage and less and later in the late stage, resulting in a slightly higher door-opening latency in the late stage than in the early stage. Under the familiar or unfamiliar rat condition, the “continuously touch sensor” action appeared more in the late than early stage, resulting in a slightly lower door-opening latency in the late than early period. When similar rats were trapped, the rats touched the sensor continuously more frequently and appeared for the first time earlier and earlier as the number of days of testing increased, probably reflecting their increasing eagerness to help similar rats and release them from being trapped.

#### 3.2.2 Reversal of the social contact procedure in Experiment 1

In Experiment 1, to verify that the significant difference in door-opening latency between the social and non-social contact groups was indeed caused by the experimental condition of whether social contact could be made, we took another 11 Sprague-Dawley rats of the same age, all of which had learned to open the door for 15 days after the test, and reversed the procedure on the 9th day of the test, at which time the social contact group (*n =* 6) was opening the door but could not enter the trapped chamber to make contact with the trapped rats, whereas the non-social contact group (*n* = 5) could enter the trapped chamber to make contact with the trapped rats after opening the door. The reversal procedure lasted seven days, and the results are shown in **Fig. 6** and **Table S6**. Repeated measures ANOVA with experimental treatment as the between-group variable and different restrainer conditions and number of tests as the within-group variables showed that the experimental treatment main effect was not significant, *F*(1, 9) = 3.30, *p* = .103, *η*^2^ _*partial*_ = .27, that is, indicating that there was no significant difference between the door-opening latency in the social and non-social contact groups. The main effect of the restrainer conditions (empty restrainer, familiar rat) was not significant, *F*(1, 9) = 2.82, *p* = .127, *η*^2^ _*partial*_ = .24, that is, the experimental treatments did not differ in the different restrainer conditions. The experimental treatment and restrainer condition interaction were not significant, *F*(1, 9) = 0.00, *p* = .955, *η*^2^ _*partial*_ = .00; the test number main effect was significant, *F*(14, 126) = 3.46, *p <* .001, *η*^2^ _*partial*_ = .28; test number and experimental treatment interaction were significant, *F*(14, 126) = 12.34, *p* < .001, *η*^2^ _*partial*_ = .58; test number and restrainer condition interaction were not significant, *F*(14, 126) = 0.88, *p <* .583, *η*^2^ _*partial*_ = .09; and the number of tests, restrainer condition, and experimental treatment interaction were significant, *F*(14, 126) = 2.33, p = .007, *η*^2^ _*partial*_ = .21. Further testing revealed that the difference in door-opening latency between the social and non-social contact groups was not significant on Days 1, 2, 6, and 8 (*p =* .092; *p =* .309; *p* = .500; *p* = .289). On Days 3, 4, 5, and 7, the difference in door-opening latency between the social and non-social contact groups was significant (*p* < .001; *p* < .001; *p* = .007; *p* = .017), that is, the door-opening latency in the socially exposed group was shorter than that of the non-socially exposed group. On Days 9, 12, and 14, there was no significant difference in door-opening latency between the social and non-social contact groups (*p =* .239; *p =* .228; *p =* .072). On Days 10, 11, 13, and 15, there was a significant difference in door-opening latency between the social and non-social contact groups (*p* = .029; *p* = .031; *p* = .032; *p* = .009), that is, the socially exposed group had a longer door-opening latency than did the non-socially exposed group (after reversal).

**Figure 6.**
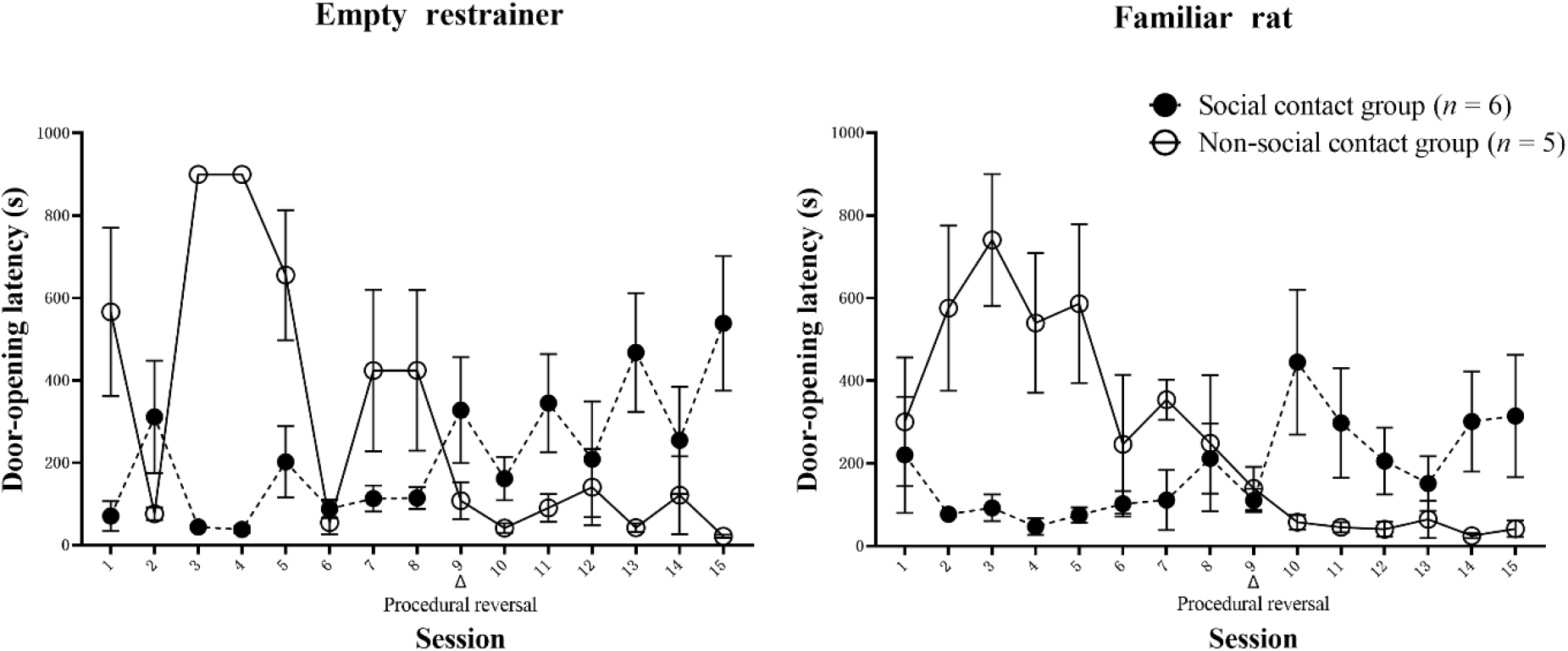
Door-opening latency for the social and the non-social contact groups in the case of procedural reversal

#### 3.2.3 Experiment 2: Effects of social contact on helping behavior in the escapable condition

The helping behavior test in Experiment 2 was conducted over 9 days, as in Experiment 1, and was divided into the early-, middle-, and late-test periods according to the number of days tested (**Fig. 7**). The results of the ANOVA on door-opening latency (**Table S7-S8**) showed that the main effect of the experimental treatment was significant, *F*(1, 26) = 123.75, *p* < .001, *η*^2^ _*partial*_ = .83, and the door-opening latency was shorter in the social (76.6±7.9 s) than in the non-social contact group (405.7±28.5 s), which supports hypothesis (2) and indicates that social contact are an important motivation for the maintenance of helping behavior in rats, and the free rats had a shorter door-opening latency in contexts in which they were able to socially engage with the trapped rats.

**Figure 7.**
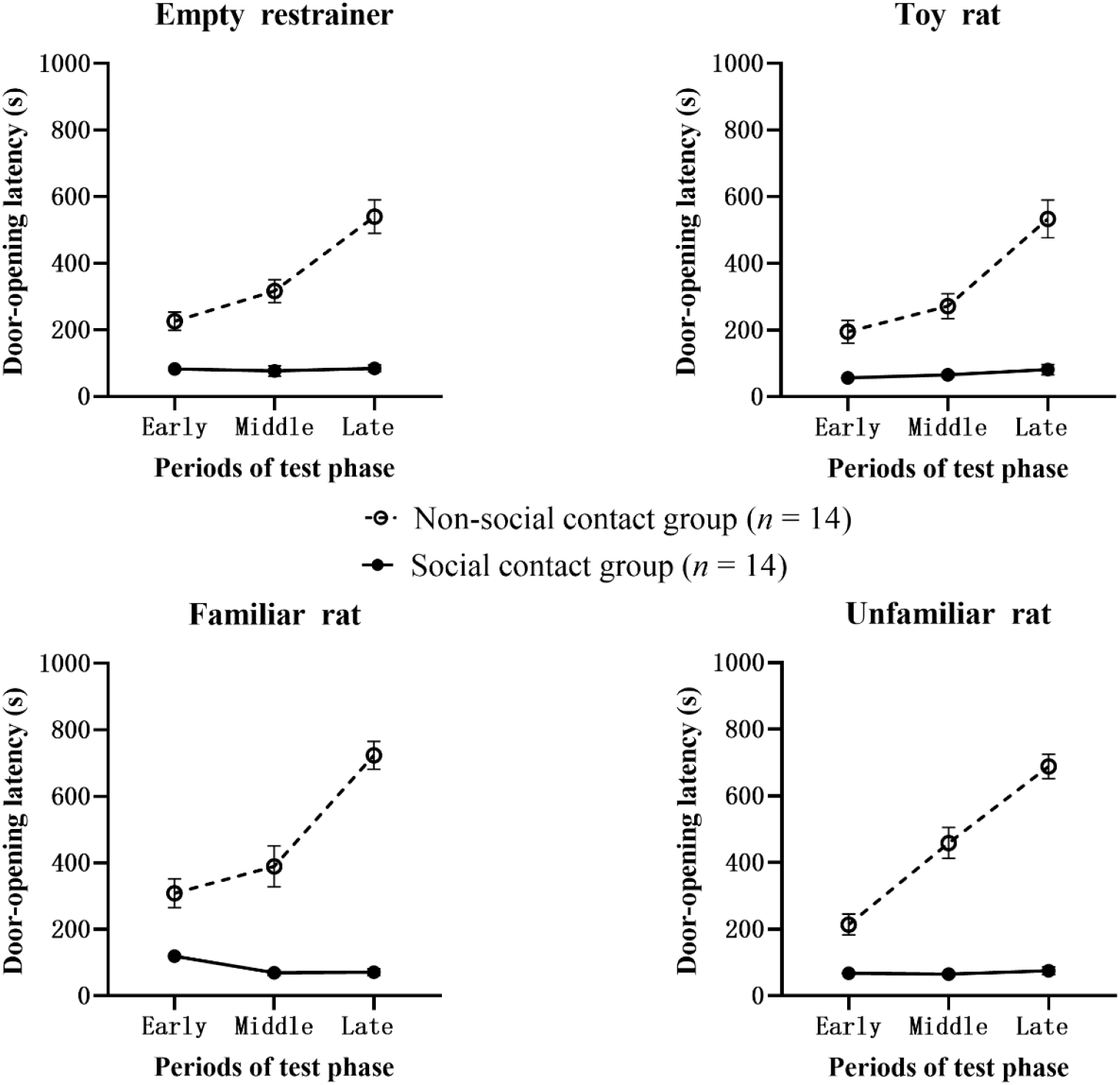
Door-opening latency in the test phase in Experiment 2

The main effect for the test period was significant, *F*(2, 52) = 77.10, *p <* .001, *η*^2^ _*partial*_ = .75, and the door-opening latency was significantly shorter in the early-test period (158.9±19.9 s) than in the middle- (214.4±32.8 s) and late-test periods (350.1±56.0 s), whereas the interaction between the test period and experimental treatment was also significant, *F*(2, 52) = 77.91, *p* < .001, *η*^2^ _*partial*_ = .75, as while the social contact group maintained a short door-opening latency throughout the early- (81.9±7.4s), middle- (69.5±7.9 s) and late- (78.3±11.2 s) test periods, the non-social contact group had a significantly shorter door-opening latency in the early- (235.9±25.9 s) than in the middle- (359.3±34.3 s) and late-test (621.9±39.2 s) periods, which again support hypothesis (2) and suggest that the door-opening latency will become longer and longer when social contact with trapped rats is not possible. The main effect of the restrainer condition was significant, *F*(3, 78) = 9.97, *p* < .001, *η*^2^ _*partial*_ = .28, with the door-opening latency in the object condition (211.3±30.5 s) shorter than that in the conspecific condition (271.0±40.8 s), which does not support hypothesis (1). The difference in door-opening latency between the familiar and unfamiliar rat conditions was not significant, *t*(26) = 1.39, *p* = 0.517, which does not support hypothesis (5).

#### 3.2.4 Comparative analysis: Experiments 1 and 2

Comparing the door-opening latency between the social contact groups in Experiments 1 and 2, a repeated measures ANOVA with the study (Experiments 1 and 2) as the between-group variable and the condition and test period as the within-group variables (**Table S9-S10**) showed a non-significant main effect, *F*(1, 26) = 2.27, *p =* .144, *η*^2^ _*partial*_ = .08. Comparing the door-opening latency between the non-social contact groups’ door-opening latency (**Table S11-S12**) in Experiments 1 and 2, the results showed a significant main effect, *F* (1, 26) = 6.09, *p =* .021, *η*^2^ _*partial*_ = .19, and a significantly shorter door-opening latency in Experiment 2 (405.7±28.5 s) than in Experiment 1 (522.8±38.0 s). As the only difference between Experiments 2 and 1 was the accessibility of the dark chamber, the results supported hypothesis (3), indicating that the presence of the dark chamber helped alleviate their own distress, resulting in a shorter door-opening latency and increased continuous door-opening behavior in the non-social contact group.

#### 3.2.5 Experiment 3: Effect of prior distress experience and social contact on helping behavior in escapable conditions

The helping behavior test in Experiment 3 was conducted over six days and was divided into early-, middle-, and late-test periods (**Fig. 8**). The analysis of door-opening latency (**Table S13-S14**) showed that the main effect of the experimental treatment was significant, *F*(1, 24) = 28.50, *p* < .001, *η*^2^ _*partial*_ = .54, that is, the door-opening latency was shorter in the social contact group (108.9±13.0 s) than in the non-social contact group (391.3±52.5 s), which supports hypothesis (2) and suggests that social contact is the maintenance of helping behavior in rats. The main effect of the test period was significant, *F*(2, 48) = 11.24, *p <* .001, *η*^2^ _*partial*_ = .32, and the door-opening latency was significantly shorter in the early-test period (181.4±27.6 s) than in the middle- (269.6±48.3 s) and late-test periods (299.3±48.6 s). The interaction between the test period and experimental treatment was significant, *F*(2, 48) = 19.12, *p* < .001, *η*^2^ _*partial*_ = .44. Additional tests revealed that the door-opening latency in the non-social contact group was significantly shorter in the early- (234.1±47.9 s) than in the middle- (432.4±73.0 s) and late-test (507.4±55.3 s) periods regardless of the social contact experience, again supporting hypothesis (2), suggesting that when social contact was not available, the non-social contact group’s door-opening latency increased gradually as the experimental schedule progressed.

**Figure 8.**
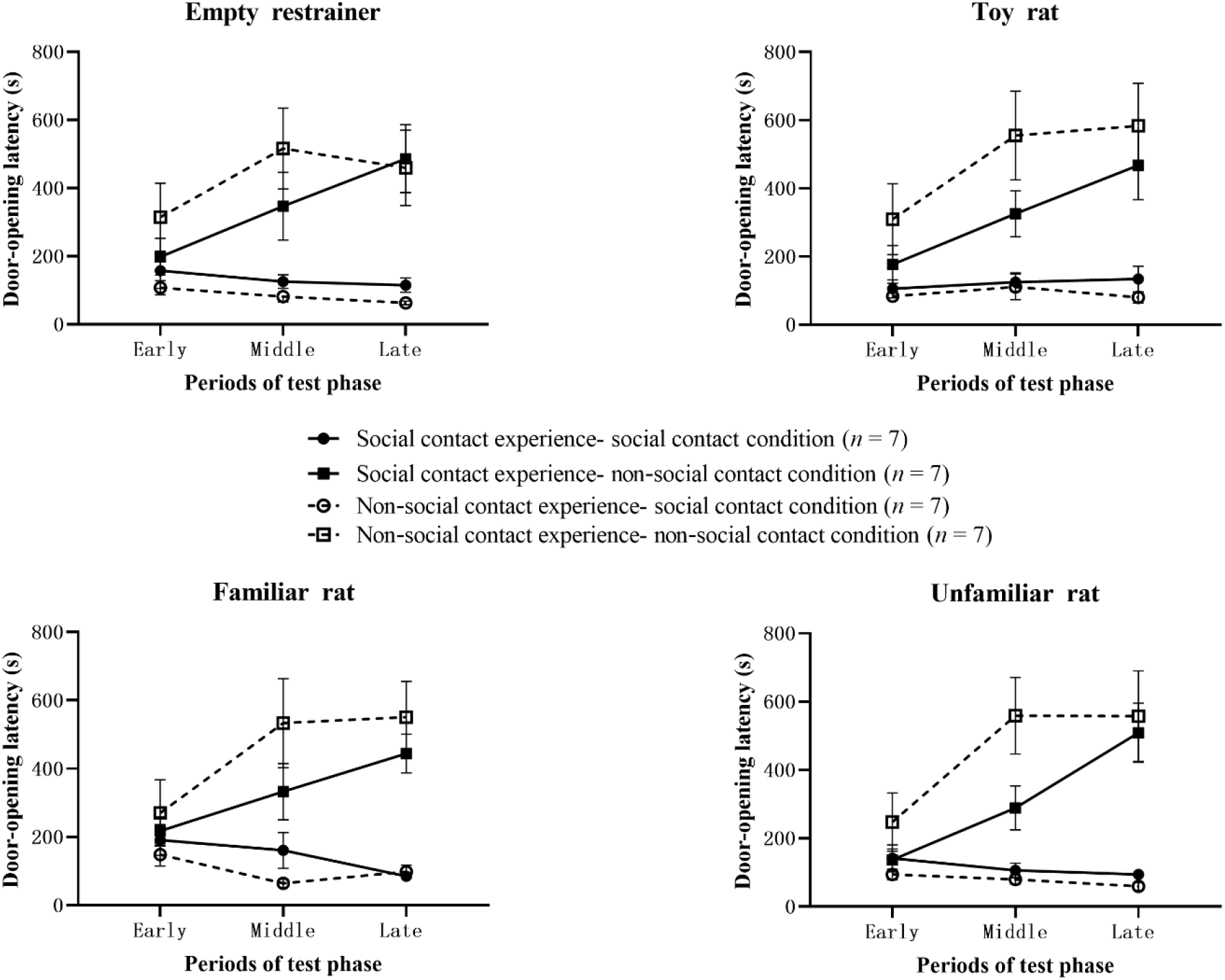
Door-opening latency in the test phase in Experiment 3

The main effect of social contact experience was not significant, *F*(1, 24) = 0.69, *p* = .414, *η*^2^ _*partial =*_ .03, and the interaction between social contact experience and experimental treatment was not significant, *F*(1, 24) = 2.47, *p* = .129, *η*^2^ _*partial*_ = .09. However, multiple comparisons revealed that the door-opening latency in rats with prior social contact experience (228.1±36.0 s) was shorter than that in rats without prior social contact experience (272.1±68.0 s); in the non-social contact group, the door-opening latency was shorter in rats with prior social contact experience group (327.7±42.4 s) than in the non-social contact experience group (454.9±93.7 s), indicating that social contact experience would have an effect on helping behavior in rats, partially supporting hypothesis (4). The main effect of the restrainer condition was not significant, *F*(3, 72) = 0.44, *p =* .727, *η*^2^ _*partial*_ = .02, and the door-opening latency in the object condition (251.3±38.2 s) was barely different from that in the similar condition (248.9±38.4 s), which does not support hypothesis (1). The difference between the door-opening latency in the familiar and unfamiliar rat conditions was not significant, *t*(24) = 1.59, *p* = 0.404, which does not support hypothesis (5).

#### 3.2.6 Comparative analysis: Experiments 2 and 3

To maintain temporal consistency, data from the first six days of Experiment 2 were taken and averaged every two days according to the number of days tested and divided into early-, middle-, and late-periods. Comparing the differences in door-opening latency between the social contact groups in Experiments 2 and 3, a repeated measures ANOVA was conducted with the study as the between-group variable and the condition and test period as the within-group variables (**Table S15-S16**), showing that the main effect was significant, *F*(1, 26) = 5.08, *p* = .033, *η*^2^ _*partial*_ = .16, but the door-opening latency was longer in the rats in Experiment 3 (108.9±13.0 s) than in Experiment 2 (75.7±7.0 s). Similarly, comparing the door-opening latency between the non-social contact groups in Experiments 2 and 3 (**Table S17-S18**) showed that the main effect was not significant, *F*(1, 26) = 2.48, *p* = .128, *η*^2^ _*partial*_ = .09, and that the door-opening latency was longer in rats in Experiment 3 (391.3±52.5 s) than in Experiment 2 (297.6±28.1 s). As Experiments 2 and 3 differed only in the role transformation of the subjects, the comparison revealed that the door-opening latency was longer in Experiment 3 than in Experiment 2 for the socially and non-socially exposed groups, that is, past trapping experience did not cause the rats’ helping behavior to unfold more quickly, which was inconsistent with the hypothesis that past trapping experience were relevant in hypothesis (4).

## 4 Discussion

### 4.1 Empathy may not be the main cause for helping behavior

Although empathy is often considered a basic ability of social cognition, the comparative cognition perspective suggests that empathy comprises emotional (a bottom-up automatic process driven by stimuli, which is a substitute experience of other individuals’ emotional states), cognitive (the process by which individuals identify and understand others’ emotional feelings and their causes based on certain conceptual systems and rules), and behavioral (the process by which individuals show behavioral tendencies or reactions in the process of empathy) empathy (Wang et al., 2021; Wei & Su, 2019). The “Matryoshka doll” model of empathy evolution (de Waal & Preston, 2017) suggests that the core of empathy is the perception-action mechanism, which means that when the observer perceives the state of the other person, the nervous system will spontaneously activate individual representations pertaining to that state or situation, thus enabling the observer to understand the other person’s emotional state or to trigger similar emotional states of their own. Bartal et al. (2011), Cox and Reichel (2020), and Sato et al. (2015) showed that rats continued to engage in helping behaviors even when social contact was not possible, thus determining that other than empathic motivation, there seems to be no other possible plausible explanation (Mogil, 2012). In contrast, although behaviors that characterize empathy, such as sniffing and rigidity, were observed in free rats toward trapped ones, they were not sufficient to suggest that empathy was the primary motivation for maintaining helping behavior in rats. First, the body writhing and twisting and increased excretion of the trapped rats restrained in the restrainer reflected that they were in a state of intense stress and needed help. Second, rats can identify conspecifics through olfactory cues, obtain information on social objects (Brown, 1979), and determine the stressful, anxious state of conspecifics (Galef & Wigmore, 1983), and identify whether the other needs help (Gerber et al., 2020; Schneeberger et al, 2020). Free rats in the non-social contact group were accompanied by a great extent of rigidity in trials where the door was opened very slowly or even not, and rigidity is an important indicator of emotional contagion and empathy in rats (Conti et al., 2012). Behaviors such as sniffing and rigidity in free-ranging rats suggest that they acquire information from their conspecifics and that emotional contagion occurs. However, compared to the object condition, free-ranging rats do not open the door more quickly because their conspecifics are in distress. This study found, through qualitative coding, that when there were conspecifics in the restrainer (familiar and unfamiliar rat conditions), the first action the free rats engaged in while entering the middle chamber was sniffing the holes, whereas when the restrainer condition was the object (empty restrainer and toy rat conditions), the free rats usually walked around first upon entering the middle chamber. Late in the test, the non-social contact group did not help open the door, but remained frequently at the sniffing hole closest to the trapped rat. Thus, the free rats were able to share the emotional state of the trapped rats regardless of whether social contact was available or not. This suggests that (emotional) empathy is not the main reason for maintaining helping behavior in rats, and that empathy, which includes emotional, cognitive, and behavioral components, is a theoretical description of helping behavior.

### 4.2 Helping behavior may be an adjunct to the pursuit of social contact and environmental “fun” in rats

The results showed that while controlling for the ability of free rats to socially engage with trapped rats, there was a significant difference in door-opening latency between the socially and non-socially engaged groups of free rats, supporting the social engagement hypothesis. Helping behavior is more likely to occur when rats are able to socially engage with their own kind (Heslin & Brown, 2021; Schwartz et al., 2017; Silberberg et al., 2014), and the inability to socially engage will result in a gradual decrease in door-opening behavior in free rats. If the social and the non-social contact group conditions in Experiment 1 were reversed (i.e., at this point the social contact group could not enter the trapped chamber after opening the door, whereas the non-social contact group could enter the trapped chamber after opening the door), the door-opening latency results were similarly reversed. These results are consistent with the findings of Schwartz et al. (2017) and Silberberg et al. (2014). A model of pain empathic behavior in rats pioneered by domestic researchers suggests the presence of empathic comfort behaviors in rats driven by social interactions with trapped rats in pain (C.-L. Li et al., 2018; Z. Li et al., 2014). Fighting and playing are common forms of games in the species (Henry & Herrero, 1974; Kight et al., 2021; Pellis & Pellis, 1997), from which social contact can activate, and reinforce, reward-related circuits. Social play during adolescence is significant for mammalian growth and development (Graham & Burghardt, 2010; Michael & Székely, 2019), The subjects in this study were adolescent rats, whose social interactions with their littermates are of high value (Douglas et al., 2004; Latané& Werner, 1978; Vanderschuren et al., 1997). Thus, helping behavior is more likely to occur when free rats in the social contact group can interact and play with trapped (familiar cagemates, unfamiliar rats) or toy rats after opening the door.

In the experimental setup (**Fig. 1**), the rectangular platform at the end of the restrainer added interest to the trapped chamber as did the restrainer that could be drilled into after opening the door. A random video tally of one day (Day 5 of Experiment 2) showed that nearly 80% of free rats climbed onto the platform or burrowed into the restrainer in the empty restrainer and toy rat conditions, suggesting that the restrainer in the trap room served as a “fun facility” to facilitate the opening of the door by free rats, which is consistent with previous studies (Bartal et al., 2011; Ueno et al., 2018). Ueno et al. (2018) suggested that the emergence of rescue-like behavior in rodents may be the result of interest in the restrainer, rather than empathy. They placed two restrainers in their experiment, in which one held a cage mate and the other was empty, and left the rat to choose freely, resulting in rats showing more attention to the empty restrainer and preferentially staying in it instead of rescuing the trapped cage mate. Silberberg et al. (2014) found that rats had a shorter delay in opening the door in the first five days, possibly because the motivation to explore the environment while entering a new space promoted them to open the door sooner, and helping behavior is an accessory to exploring the environment. Thus, changes in the environment have a moderating effect on rats’ motivation for helping behavior, and when social exposure is more interesting, helping behavior will not be a purely prosocial behavioral motivation, but rather a way for rats to gain social contact or explore the interesting environment.

### 4.3 The presence of a dark chamber reduces pain and helps promote helping behavior in rats

Comparing the results of Experiments 1 and 2, we found that when the dark chamber was accessible, there was an increase in door-opening latency for free rats rescuing trapped rats by touching the sensor, which is consistent with previous findings (Carvalheiro et al., 2019). Bartal et al. (2016) showed that administering anxiolytic drugs to free rats significantly reduced door-opening behavior, suggesting that the pain felt by free rats is necessary to open the restrainer door. Instead of using anxiolytic drugs in this study, the free rats were given the option to escape to a dark chamber to alleviate their pain. The comparison revealed that the difference in door-opening latency between Experiments 1 and 2 without the social contact group was significant, and that the door-opening latency was significantly longer in Experiment 1 than 2. As there was no difference in Experiment 2 except for the presence of more dark chambers than in Experiment 1, this difference may have resulted from the presence of dark chambers. In Experiment 1, when free rats were unable to alleviate their pain by hiding in the dark chamber, the free rats in the non-social contact group began to show more non-opening behavior on Day 3, whereas in Experiment 2 the non-opening behavior of the free rats in the non-social contact group began to appear around Day 6. Similarly, on the last day, 73 % of the rats in Experiment 1 did not open the door, and 50 % of the free rats in Experiment 2 did not open the door. This suggests that the presence of the dark chamber increased the continuous door-opening behavior of the free rats in the non-social contact group. In the social contact group, the door-opening latency was slightly higher in Experiment 2 than 1. The video of the dark chamber in the social contact group showed that the free rats did not show grooming or rigid behavior while entering the dark chamber. They mostly shuttled between the light and dark chambers or explored the dark chamber walls, suggesting that the presence of the dark chamber increased the exploration area in the social contact group. For the non-social contact group, the presence of the dark chamber made the free-rangers less distressed and helped them maintain continuous door-opening behavior longer, whereas for the social contact group, the presence of the dark chamber diverted the free-rangers’ attention.

We used other indicators to explore the effect of perceived distress on helping behaviors in free rats. First, we defined behaviors that indicate anxiety or distress in free rats. The time spent in the dark chamber was measured, which is one of the indicators of the emotional state of the rats (Bailey & Crawley, 2009; Bourin & Hascoët, 2003), and longer time spent in the dark chamber indicated that they were more anxious or distressed. The duration the free rats stayed in the dark chamber before the door was opened in Experiment 2 (**Table S19**) revealed that the non-socially exposed group was in the dark chamber four times longer than was the socially exposed group, suggesting that both groups needed to alleviate their distress, but that the non-socially exposed group had stronger motivation to alleviate its distress and that relief helped promote helping behavior. Although the free rats did not open the door, they spent most of their time in the middle chamber rather than the dark chamber, suggesting that the motivation to alleviate their pain was not stronger than that for social contact.

### 4.4 Social contact experience rather than trapping experience promote helping behavior in rats

Sato et al. (2015) demonstrated that after role reversal, previously submerged rats opened the door significantly faster than did previously non-submerged rats, suggesting that previous trapping experience accelerated their speed in opening the door to rescue their partners. The cooperative behavior of female rats was influenced by the prior experience of being helped, independent of the identity of the helper (Rutte & Taborsky, 2007). In Experiment 3, role reversal was used to investigate whether trapping experience could influence helping behavior, and it was found that past trapping experience did not result in shorter door-opening latencies in rats. Experiment III explored the effect of past social contact experience on helping behavior and found that rats with prior social contact experience had shorter door-opening latencies than did rats without social contact experience even though they were unable to interact socially after opening the door. Rats that were able to interact socially in the test had shorter door-opening latencies regardless of prior social contact experience. Templer et al. (2018) used a novel socialization test and found that rats grown in a non-social environment were more willing to approach unfamiliar rats. Rats in the non-social contact group in this study had not been exposed to other rats in previous experimental contexts, so they were strongly motivated to perform helping behaviors when they were able to engage in social contact. Thus, helping behavior was influenced by prior social contact experience more than by prior trapping experience.

### 4.5 Familiarity has a small effect on helping behavior in rats

Prosocial behavior is modulated by familiarity bias, where individuals consider who to help first based on the proximity of the relationship (Bartal et al., 2014), and inter-individual familiarity is an important determinant of how one responds to the plight of others (Meyza et al., 2017). To investigate whether familiarity modulates helping behavior in rats, we compared the door-opening latencies of adolescent rats to familiar and unfamiliar trapped rats, and found that there was no significant difference in door-opening latencies between rats regardless of familiarity.

Rodents can perceive the state of their conspecifics through sight and smell (Langford et al., 2006). Rogers-Carter et al. (2018) observed that male and female rats are socially avoidant of unfamiliar conspecifics in a stressful state, but that females show a social preference for familiar cagemates in a stressful state. Burkett et al. (2016) found that for monogamous prairie voles, prosocial behavior was only expressed in familiar companions. Rats were observed to show more grooming behavior toward their cagemates when they saw them in pain (Lu et al., 2018). Bartal et al. (2014) showed that helping behavior in rats occurs not only in familiar cagemates but also in unfamiliar conspecifics, to which they argued that rats are gregarious animals that live together based on social experience, and that they obtain sensory cues from their peers and distinguish unfamiliar conspecifics that share cues with their peers from outgroups that do not share cues with their peers through these information pathways, leading to helping behavior among familiar cagemates and genetically similar unfamiliar conspecifics. Similarly, the present study found a smaller effect of familiarity on helping behavior in rats, which may have been so because the subjects had learned to open the door in the learning phase, and their curiosity and habituation to the trapped room were the immediate triggers and default behavioral patterns of their helping behavior.

### 4.6 Helping behavior is not necessarily prosocial from the perspective of behavioral motivation

Bartal et al. (2011) found that the door-opening latency of rats showed a decreasing trend only in the presence of trapped rats, but not in other conditions. Our findings are similar, suggesting that the occurrence of helping behavior in the presence of trapped conspecifics is associated with concern for conspecifics and is prosocial in nature. In the empty restrainer and toy rat conditions, although the rats showed the behavioral outcome of “opening the door,” the motivation for the behavior was the fun of the situation (or the size of the space to explore) and was not prosocial. Prosocial behavior refers to behaviors that individuals voluntarily engage in that are beneficial to others and society, including sharing, donating, comforting, cooperating, helping, and self-sacrificing (Snippe et al., 2018). In contrast, Ryan et al. (1989) argued that prosocial behaviors are motivated by certain internal or external motives, such as seeking rewards, avoiding punishment, and gaining social reputation, which make it necessary for individuals to engage in prosocial behaviors. Therefore, in terms of behavioral outcomes, helping behavior is beneficial to others and the society, and is a type of prosocial behavior. However, from the perspective of behavioral motivation, helping behavior is not necessarily prosocial.

### 4.7 Cross-species comparison of motivations for helping behavior

This study illustrated that the desire for social contact and contextual fun are strong drivers of helping behavior in rodents. The search for social contact outweighs the motivation to alleviate personal distress, whereas empathy may not be the main reason for maintaining helping behavior in rats. Such motivations for helping behavior are present in rodents and nonhuman primates and human infants. Nonhuman primates tend to socially engage (Harlow & Zimmerman., 1959), care for their own kind and offer help spontaneously (de Waal, 2007; Horner et al., 2011) or comfort to alleviate their own distress in a situation (Koski & Sterck, 2010). Social interactions are a strong motivating factor driving helping behavior in children (Giner Torréns et al., 2021), who genuinely care about the well-being of others and show it through changes in their pupils (Hepach et al., 2012), whereas the distressed state of others causes infants to feel pain, tension, and sadness (Michael & Székely, 2019). We suggest that there may be a circling model for the development of motivation for helping behavior.

Mammals have the intrinsic ability to empathize with the pain and pleasure of their surrounding kind through primitive emotional contagion (Panksepp, 2011), which is predicated on the development of the ability to distinguish between self-states and others (Hoffman, 1998). Where help is necessary, individuals spontaneously process information, decode similar emotions, achieve state matching, and generate emotional empathy according to perceptual-motor mechanisms (de Waal & Preston, 2017), which is the process underlying the occurrence of helping behaviors. However, the ability to experience others with emotional contagion does not necessarily lead to helping behaviors. Emotional empathy can lead to motivations for helping behaviors that are other-centered and sincerely concerned for the well-being of others, or to motivations for helping behaviors that are designed to alleviate one’s own suffering owing to excessive emotional arousal into one’s own suffering. Anticipated rewards motivate helping behavior, and social interactions and contact have reward value, accompanied by the transmission of neuropeptide signals such as oxytocin and dopamine, and the activation of brain regions such as the ventral striatum and medial prefrontal cortex (Marsh et al., 2014), leading to positive emotional experience.

Thus, the occurrence and acquisition of motivation to help in helping behavior situations involving emotional empathy, to alleviate one’s own suffering, and for social contact, are common between humans and animals. However, humans have more advanced cognition and may also help comply with social norms (Siposova et al., 2021), gain social acceptance (Oarga et al., 2015), and achieve self-worth, among other things. There may be a similar evolutionary developmental sequence of individuals’ motivations for helping behavior from emotional-behavioral to cognitive systems and from lower to higher levels.

## 5 Conclusion

The following conclusions were drawn from this study. (1) The possibility and experience of social contact and the fun nature of the situation are strong driving forces of helping behavior in rodents; (2) relieving one’s pain can facilitate helping behavior in rodents; and (3) from a comparative cognitive perspective, empathy may not be the main reason for maintaining helping behavior in rodents, but may be a theoretical description of their helping behavior process.

## Data Availability

All data generated to support the findings of this study are available at https://doi.org/10.57760/sciencedb.o00115.00014.

## Author contributions

S Han: Conceptualization, Investigation, Methodology, Formal analysis, Visualization, Writing— original draft and editing the manuscript; YQ Chen: Conceptualization, Investigation, Methodology, Formal analysis, Visualization, Writing—original draft and editing the manuscript; B Zheng: Investigation; YX Wang: Investigation; B Yin: Conceptualization, Supervision, Methodology, Writing—review & editing, Funding acquisition

## Competing interests

The authors declare that the research was conducted in the absence of any commercial or financial relationships that could be construed as a potential competing interest.

## Additional information

### Supplementary Information

The supplementary figures and tables are available at a separate file.

**Correspondence** and requests for additional materials should be addressed to B.Y.

## Figure Legends

**Figure 1**: At the beginning of the experiment, the trapped rats were placed in the restrainer. The restrainer door and door 1 were closed. The free rat was placed in the middle chamber. In Experiment 1, when door 2 was closed, the free rats in the social contact group could move in the middle chamber and the trapped chamber after touching the sensor to open the door, whereas the free rats in the non-social contact group could move only in the middle chamber before and after touching the sensor. In Experiment 2, door 2 was open. During the experiment, all the free rats could move in the middle and dark chamber, and the free rats in the social contact group could move in the trapped chamber after opening door 1. The range of activity of the free rat in Experiment 3 was the same as that in Experiment 2.

**Figure 2**: The horizontal coordinate represents the number of learning sessions (days). In this study, it was 16 sessions (days). First, Days 1 to 10 constitute learning stage 1, where the subject needs to touch the sensor once to open the door; Days 11 to 13 constitute learning stage 2, where the subject needs to touch the sensor 5 times in a row (fixed Ratio - 5 times, FR - 5) to open the door; and Days 14 to 16 constitute learning stage 3, where the subject needs to not touch the sensor 5 times in a row to open the door. The results are expressed as ***M ± SE***, hereinafter.

**Figure 3-8**: No legends necessary.

## Notes

https://doi.org/10.57760/sciencedb.o00115.00014

### Competing Interest Statement

The authors have declared no competing interest.

### Summary of Updates

The decimal notation for all inferential statistical results in the main text and attached tables has been modified to conform to standard usage. Specifically, the leading zeros have been removed from the decimal points of the P-values and partial eta-squared values. Additionally, errors in the calculation of standard errors (SE) for some descriptive statistics results in both the main text and attached tables have been corrected. The degrees of freedom (df) values for some inferential statistical results in the main text that were found to be inaccurate have also been corrected. Finally, modifications have been made to the notation for t-test results, including the inclusion of the df values.

## References

Bailey, K. R., & Crawley, J. N. (2009). Chapter 5: Anxiety-related behaviors in mice. In J. J. Buccafusco (Ed.) Methods of Behavior Analysis in Neuroscience (pp. 77–101). CRC Press.

Bamberger, P. A., Geller, D., & Doveh, E. (2017). Assisting upon entry: Helping type and approach as moderators of how role conflict affects newcomer resource drain. Journal of Applied Psychology, 102(12).

Bartal, I. B.-A., Breton, J. M., Sheng, H., Long, K. L., Chen, S., Halliday, A., Kenney, J. W., Wheeler, A. L., Frankland, P., Shilyansky, C., & others. (2021). Neural correlates of ingroup bias for prosociality in rats. Elife, 10.

Bartal, I. B.-A., Decety, J., & Mason, P. (2011). Empathy and prosocial behavior in rats. Science, 334(6061), 1427–1430.

Bartal, I. B.-A., Rodgers, D. A., Sarria, M. S. B., Decety, J., & Mason, P. (2014). Prosocial behavior in rats is modulated by social experience. Elife, 3, e01385.

Bartal, I. B.-A., Shan, H., Molasky, N. M., Murray, T. M., Williams, J. Z., Decety, J., & Mason, P. (2016). Anxiolytic treatment impairs helping behavior in rats. Frontiers in Psychology, 7, 850.

Bibb, L., James, M., Virginia, J., Bernard, L., & Howard, C. (1972). Stimulus determinants of social attraction in rats. Journal of Comparative and Physiological Psychology, 79(1), 13.

Bourin, M., & Hascoët, M. (2003). The mouse light/dark box test. European Journal of Pharmacology, 463(1–3), 55–65.

Brown, R. E. (1979). Mammalian Social Odors: A Critical Review. In J. S. Rosenblatt, R. A. Hinde, C. Beer, & M.-C. Busnel (Eds.), Advances in the Study of Behavior (pp. 103–162). Academic Press. https://doi.org/10.1016/S0065-3454(08)60094-7

Burkett, J. P., Andari, E., Johnson, Z. V., Curry, D. C., de Waal, F. B., & Young, L. J. (2016). Oxytocin-dependent consolation behavior in rodents. Science, 351(6271), 375–378.

Campos, A.C., Ferreira, F.R., Guimarães, F.S., & Lemos. J.I. (2010). Facilitation of endocannabinoid effects in the ventral hippocampus modulates anxiety-like behaviors depending on previous stress experience. Neuroscience, 2. https://doi.org/10.1016/j.neuroscience.2010.01.062.

Carvalheiro, J., Seara-Cardoso, A., Mesquita, A. R., de Sousa, L., Oliveira, P., Summavielle, T., & Magalhães, A. (2019). Helping behavior in rats (Rattus norvegicus) when an escape alternative is present. Journal of Comparative Psychology, 133(4), 452.

Clay, Z., & de Waal, F. B. (2013a). Bonobos respond to distress in others: Consolation across the age spectrum. PLoS ONE, 8(1), e55206.

Clay, Z., & de Waal, F. B. (2013b). Development of socio-emotional competence in bonobos. Proceedings of the National Academy of Sciences, 110(45), 18121–18126.

Conti, G., Hansman, C., Heckman, J. J., Novak, M. F., Ruggiero, A., & Suomi, S. J. (2012). Primate evidence on the late health effects of early-life adversity. Proceedings of the National Academy of Sciences, 109(23), 8866–8871.

Costa, R., Tamascia, M. L., Nogueira, M. D., Casarini, D. E., & Marcondes, F. K. (2012). Handling of adolescent rats improves learning and memory and decreases anxiety. Journal of the American Association for Laboratory Animal Science, 51(5), 548–553.

Cox, S. S., & Reichel, C. M. (2020). Rats display empathic behavior independent of the opportunity for social interaction. Neuropsychopharmacology, 45(7), 1097–1104.

de Waal, F. B. M., & Preston, S. D. (2017). Mammalian empathy: Behavioural manifestations and neural basis. Nature Reviews Neuroscience, 18(8), 498–509. https://doi.org/10.1038/nrn.2017.72

de Waal, F. B. M. (2007). With a little help from a friend. PLoS Biology, 5(7), e190.

Douglas, L. A., Varlinskaya, E. I., & Spear, L. P. (2004). Rewarding properties of social interactions in adolescent and adult male and female rats: Impact of social versus isolate housing of subjects and partners. Developmental Psychobiology: The Journal of the International Society for Developmental Psychobiology, 45(3), 153–162.

Galef, B. G., & Wigmore, S. W. (1983). Transfer of information concerning distant foods: A laboratory investigation of the ‘information-centre’ hypothesis. Animal Behaviour, 31(3), 748–758. https://doi.org/10.1016/S0003-3472(83)80232-2

Gerber, N., Schweinfurth, M. K., & Taborsky, M. (2020). The smell of cooperation: Rats increase helpful behaviour when receiving odour cues of a conspecific performing a cooperative task. Proceedings of the Royal Society B: Biological Sciences, 287(1939), 20202327. https://doi.org/10.1098/rspb.2020.2327

Giner Torréns, M., Dreizler, K., & Kärtner, J. (2021). Insight into toddlers’ motivation to help: From social participants to prosocial contributors. Infant Behavior and Development, 64, 101603. https://doi.org/10.1016/j.infbeh.2021.101603.

Gonzalez-Liencres, C., Juckel, G., Tas, C., Friebe, A., & Brüne., M. (2014). Emotional contagion in mice: the role of familiarity. Behavioural Brain Research, 263, 16–21.

Graham, K. L., & Burghardt, G. M. (2010). Current perspectives on the biological study of play: Signs of progress. The Quarterly Review of Biology, 85(4), 393–418.

Hachiga, Y., Schwartz, L. P., Silberberg, A., Kearns, D. N., Gomez, M., & Slotnick, B. (2018). Does a rat free a trapped rat due to empathy or for sociality? Journal of the Experimental Analysis of Behavior, 110(2), 267–274.

Harlow, H. F., & Zimmermann, R. R. (1959). Affectional responses in the infant monkey. Science, 130(3373), 421–432.

Havlik, J. L., Vieira Sugano, Y. Y., Jacobi, M. C., Kukreja, R. R., Jacobi, J. H. C., & Mason, P. (2020). The bystander effect in rats. Science Advances, 6(28), eabb4205.

Henry, J., & Herrero, S. (1974). Social play in the American black bear: Its similarity to canid social play and an examination of its identifying characteristics. American Zoologist, 14(1), 371–389.

Hepach, R., Vaish, A., & Tomasello, M. (2012). Young Children Are Intrinsically Motivated to See Others Helped. Psychological Science, 23(9), 967–972. https://doi.org/10.1177/0956797612440571.

Hernandez-Lallement, J., Attah, A. T., Soyman, E., Pinhal, C. M., Gazzola, V., & Keysers, C. (2020). Harm to others acts as a negative reinforcer in rats. Current Biology, 30(6), 949–961.

Heslin, K. A., & Brown, M. F. (2021). No preference for prosocial helping behavior in rats with concurrent social interaction opportunities. Learning & Behavior, 49(4), 397–404.

Hiura, L. C., Tan, L., & Hackenberg, T. D. (2018). To free, or not to free: Social reinforcement effects in the social release paradigm with rats. Behavioural Processes, 152, 37–46. https://doi.org/10.1016/j.beproc.2018.03.014.

Hoffman, M. L. (1998). Varieties of empathy-based guilt. In J. Bybee (Ed.), Guilt and children. (Horner, V., Carter, J. D., Suchak, M., & de Waal, F. B. M. (2011). Spontaneous prosocial choice by chimpanzees. Proceedings of the National Academy of Sciences, 108(33), 13847–13851.

Huang L., Su Y. (2012). The lifelong development of empathy: A dual-process perspective. Psychological Development and Education, 28(4), 434–441.

Kandis, S., Ates, M., Kizildag, S., Camsari, G. B., Yuce, Z., Guvendi, G., Koc, B., Karakilic, A., Camsari, U. M., & Uysal, N. (2018). Acetaminophen (paracetamol) affects empathy-like behavior in rats: Dose-response relationship. Pharmacology Biochemistry and Behavior, 175, 146–151.

Keysers, C., Knapska, E., Moita M. A., & Gazzola V. (2022). Emotional contagion and prosocial behavior in rodents. Trends in Cognitive Sciences, 8. doi:10.1016/J.TICS.2022.05.005.

Kight, K. E., Argue, K. J., Bumgardner, J. G., Bardhi, K., Waddell, J., & McCarthy, M. M. (2021). Social behavior in prepubertal neurexin 1α deficient rats: A model of neurodevelopmental disorders. Behavioral Neuroscience, 135(6), 782.

Knapska, E., Mikosz, M., Werka, T., & Maren, S. (2010). Social modulation of learning in rats. Learning & Memory, 17(1), 35–42.

Koski, S. E., & Sterck, E. H. M. (2010). Empathic chimpanzees: A proposal of the levels of emotional and cognitiveprocessing in chimpanzee empathy. European Journal of Developmental Psychology, 7(1), 38–66. https://doi.org/10.1080/17405620902986991

Langford, D. J., Crager, S. E., Shehzad, Z., Smith, S. B., Sotocinal, S. G., Levenstadt, J. S., Chanda, M. L., Levitin, D. J., & Mogil, J. S. (2006). Social Modulation of Pain as Evidence for Empathy in Mice. Science, 312(5782), 1967–1970. https://doi.org/10.1126/science.1128322

Latané, B., & Werner, C. (1978). Regulation of social contact in laboratory rats: Time, not distance. Journal of Personality and Social Psychology, 36(10), 1128.

Lavery, J., & Foley, P. (1963). Altruism or arousal in the rat? Science, 140, 172–173.

Li, C.-L., Yu, Y., He, T., Wang, R.-R., Geng, K.-W., Du, R., Luo, W.-J., Wei, N., Wang, X.-L., Wang, Y., & others. (2018). Validating rat model of empathy for pain: Effects of pain expressions in social partners. Frontiers in Behavioral Neuroscience, 242.

Li, Z., Lu, Y.-F., Li, C.-L., Wang, Y., Sun, W., He, T., Chen, X.-F., Wang, X.-L., & Chen, J. (2014). Social interaction with a cagemate in pain facilitates subsequent spinal nociception via activation of the medial prefrontal cortex in rats. PAIN, 155(7), 1253–1261.

Lu, Y.-F., Ren, B., Ling, B.-F., Zhang, J., Xu, C., & Li, Z. (2018). Social interaction with a cagemate in pain increases allogrooming and induces pain hypersensitivity in the observer rats. Neuroscience Letters, 662, 385–388.

Márquez, C., Rennie, S. M., Costa, D. F., & Moita, M. A. (2015). Prosocial choice in rats depends on food-seeking behavior displayed by recipients. Current Biology, 25(13), 1736–1745. doi:10.1016/j.cub.2015.05.018.

Marsh, A. A., Stoycos, S. A., Brethel-Haurwitz, K. M., Robinson, P., VanMeter, J. W., & Cardinale, E. M. (2014). Neural and cognitive characteristics of extraordinary altruists. Proceedings of the National Academy of Sciences, 111(42), 15036–15041. https://doi.org/10.1073/pnas.1408440111

Mason, P. (2016). Helping Another in Distress: Lessons from Rats. The Evolution of Morality. Springer.

Mason, P. (2021). Lessons from helping behavior in rats. Current Opinion in Neurobiology. doi:10.1016/J.CONB.2021.01.001.

Meyza, K. Z., Bartal, I. B.-A., Monfils, M. H., Panksepp, J. B., & Knapska, E. (2017). The roots of empathy: Through the lens of rodent models. Neuroscience & Biobehavioral Reviews, 76, 216–234.

Michael, J., & Székely, M. (2019). Goal slippage: A mechanism for spontaneous instrumental helping in infancy? Topoi, 38(1), 173–183.

Mogil, J. S. (2012). The surprising empathic abilities of rodents. Trends in Cognitive Sciences, 3. doi:10.1016/j.tics.2011.12.012.

Oarga, C., Stavrova, O., & Fetchenhauer, D. (2015). When and why is helping others good for well-being? The role of belief in reciprocity and conformity to society’s expectations: Helping and well-being. European Journal of Social Psychology, 45(2), 242–254. https://doi.org/10.1002/ejsp.2092

Panksepp, J. (2011). Empathy and the Laws of Affect. Science, 334(6061), 1358–1359. https://doi.org/10.1126/science.1216480

Pellis, S. M., & Pellis, V. C. (1997). The prejuvenile onset of play fighting in laboratory rats (Rattus norvegicus). Developmental Psychobiology: The Journal of the International Society for Developmental Psychobiology, 31(3), 193–205.

Pitman, D.L., Ottenweller, J.E. & Natelson, B.H. (1988). Plasma corticosterone levels during repeated presentation of two intensities of restrainer stress: Chronic stress and habituation. Physiology & Behavior (1). doi:10.1016/0031-9384(88)90097-2. pp. 91–112). Academic Press. https://doi.org/10.1016/B978-012148610-5/50005-9

Rice, G. E., & Gainer, P. (1962). Altruism in the albino rat. Journal of Comparative and Physiological Psychology, 55, 123–125.

Rogers-Carter, M. M., Djerdjaj, A., Culp, A. R., Elbaz, J. A., & Christianson, J. P. (2018). Familiarity modulates social approach toward stressed conspecifics in female rats. PLoS One, 13(10), e0200971.

Rutte, C., & Taborsky, M. (2007). Generalized reciprocity in rats. PLoS Biology, 5(7), e196.

Ryan, R. M., & Connell, J. P. (1989). Perceived locus of causality and internalization: examining reasons for acting in two domains. Journal of Personality and Social Psychology, 5.

Sato, N., Tan, L., Tate, K., & Okada, M. (2015). Rats demonstrate helping behavior toward a soaked conspecific. Animal Cognition, 18(5), 1039–1047.

Schmelz, M., Grueneisen, S., Kabalak, A., Jost, J., & Tomasello, M. (2017). Chimpanzees return favors at a personal cost. Proceedings of the National Academy of Sciences, 114(28), 7462–7467.

Schneeberger, K., Röder, G., & Taborsky, M. (2020). The smell of hunger: Norway rats provision social partners based on odour cues of need. PLoS Biology, 18(3), e3000628. https://doi.org/10.1371/journal.pbio.3000628

Schwartz, L. P., Silberberg, A., Casey, A. H., Kearns, D. N., & Slotnick, B. (2017). Does a rat release a soaked conspecific due to empathy? Animal Cognition, 20(2), 299–308.

Silberberg, A., Allouch, C., Sandfort, S., Kearns, D., Karpel, H., & Slotnick, B. (2014). Desire for social contact, not empathy, may explain “rescue” behavior in rats. Animal Cognition, 17(3), 609–618.

Siposova, B., Grueneisen, S., Helming, K., Tomasello, M., & Carpenter, M. (2021). Common knowledge that help is needed increases helping behavior in children. Journal of Experimental Child Psychology, 201, 104973. https://doi.org/10.1016/j.jecp.2020.104973

Snippe, E., Jeronimus, B. F., aan het Rot, M., Bos, E. H., de Jonge, P., & Wichers, M. (2018). The Reciprocity of Prosocial Behavior and Positive Affect in Daily Life. Journal of Personality, 86(2), 139–146. https://doi.org/10.1111/jopy.12299

Stukas, A. A., & Clary, E. G. (2012). Altruism and helping behavior. Encyclopedia of Human Behavior (SecondEdition), 8, 100–107.

Templer, V. L., Wise, T. B., Dayaw, K. I. T., & Dayaw, J. N. T. (2018). Nonsocially housed rats (Ratus norvegicus) seek social interactions and social novelty more than socially housed counterparts. Journal of Comparative Psychology, 132(3), 240.

Tokuyama, N., Toda, K., Poiret, M.-L., Iyokango, B., Bakaa, B., & Ishizuka, S. (2021). Two wild female bonobos adopted infants from a different social group at Wamba. Scientific Reports, 11(1), 1–11.

Ueno, H., Suemitsu, S., Murakami, S., Kitamura, N., Wani, K., Matsumoto, Y., … & Ishihara, T. (2019). Helping-Like Behaviour in Mice Towards Conspecifics Constrained Inside Tubes.. Scientific Reports (1). doi:10.1038/s41598-019-42290-y.

Ueno, H., Suemitsu, S., Murakami, S., Kitamura, N., Wani, K., Okamoto, M., Matsumoto, Y., Aoki, S., & Ishihara, T. (2018). Empathic behavior according to the state of others in mice. Brain and Behavior, 8(7), e00986.

Vanderschuren, L. J., Niesink, R. J., & Van Pee, J. M. (1997). The neurobiology of social play behavior in rats. Neuroscience & Biobehavioral Reviews, 21(3), 309–326.

Wang Q., Liu Z., Su Y. (2021). The lifelong development of empathy and its neural basis. Scientia Sinica (Vitae), 51(06), 717–729.

Wei Q., Su Y. (2019). Empathy and its different components in preschool children. Psychology: Techniques and Applications, 7(09), 523–535.

Zahn-Waxler, C., Radke-Yarrow, M., Wagner, E., & Chapman, M. (1992). Development of concern for others. Developmental Psychology, 28(1), 126.

