## supplementary figures and tables for "An Empirical Study on the Motivation of Helping Behavior in Rats"

### Supplementary materials

#### S1. Examples of different phases of the experiment

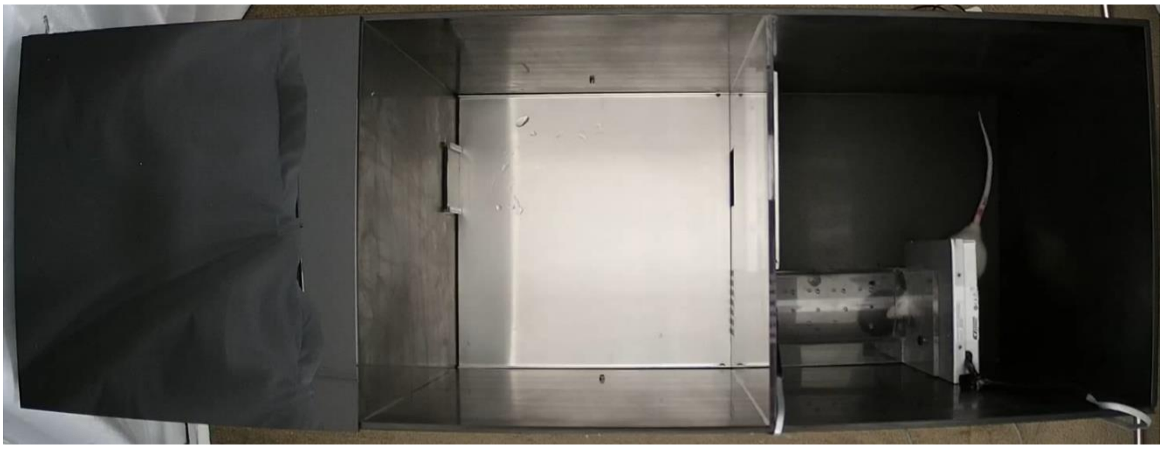

Figure S1. Experiment 1: the habituation phase for the free rat

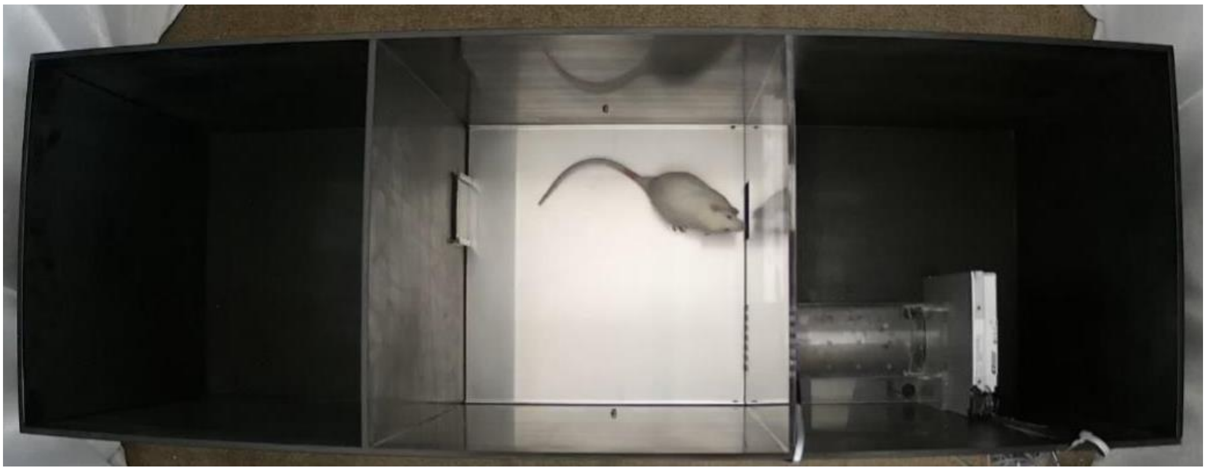

Figure S2. Experiment 1: the learning phase for the free rat

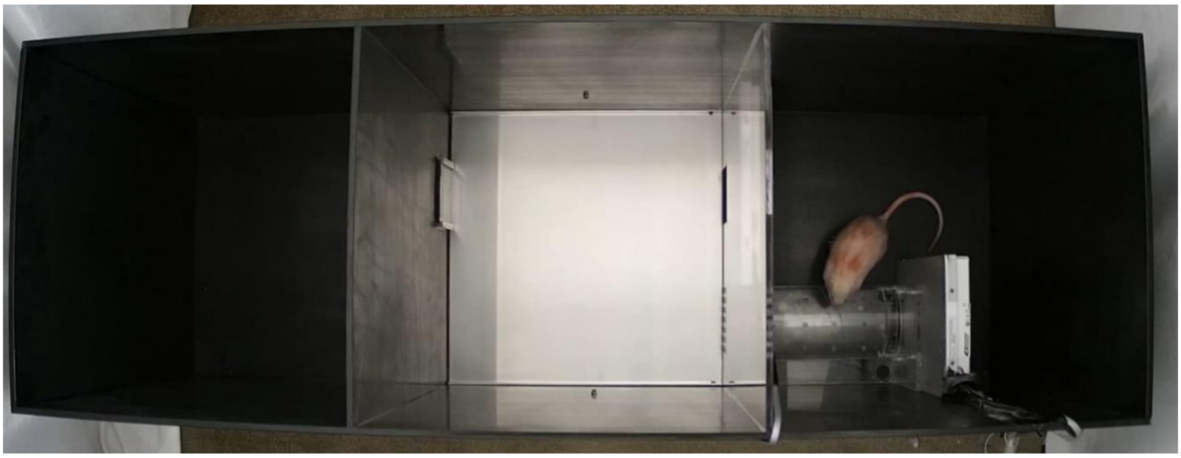

Figure S3. Experiment 1: the test phase for the free rat in the social contact group
(Empty restrainer condition)

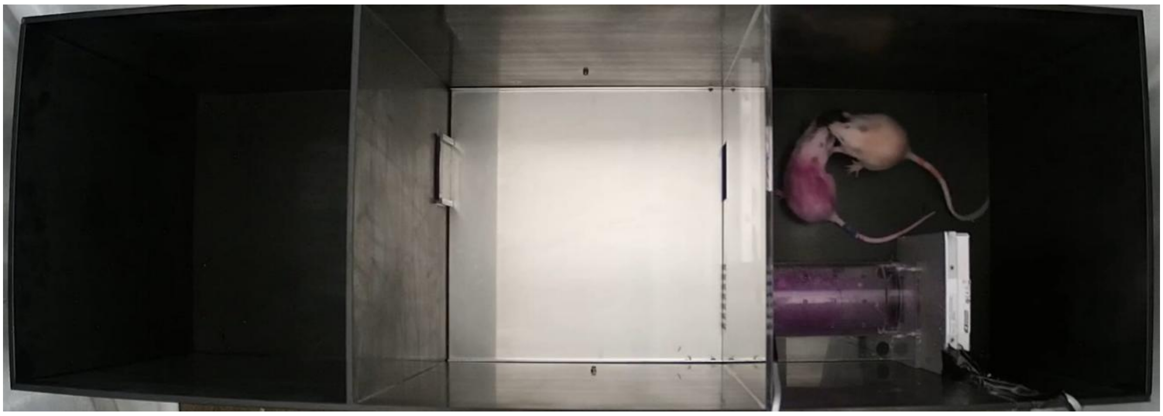

Figure S4. Experiment 1: the test phase for the free rat in the social contact group (Unfamiliar rat condition)

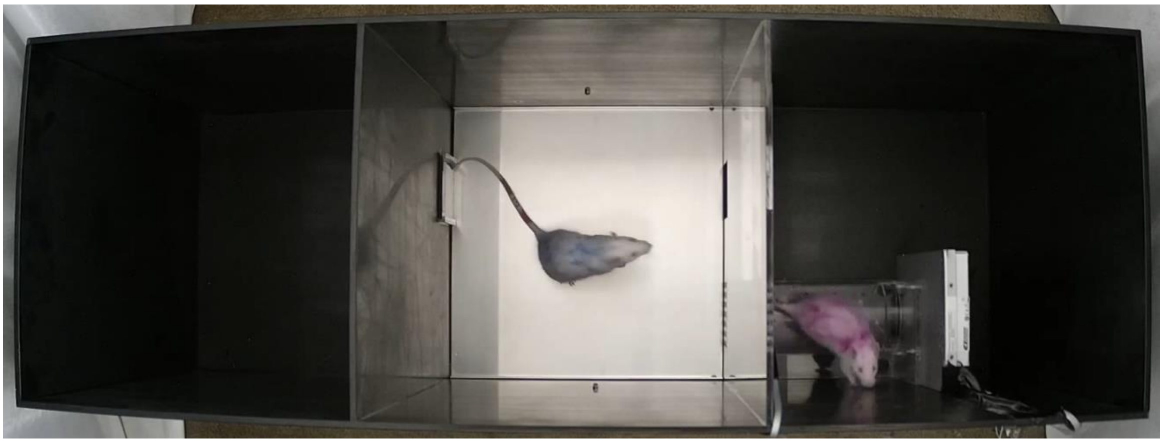

Figure S5. Experiment 1: the test phase for the free rat in the Non-social contact group (Familiar rat condition)

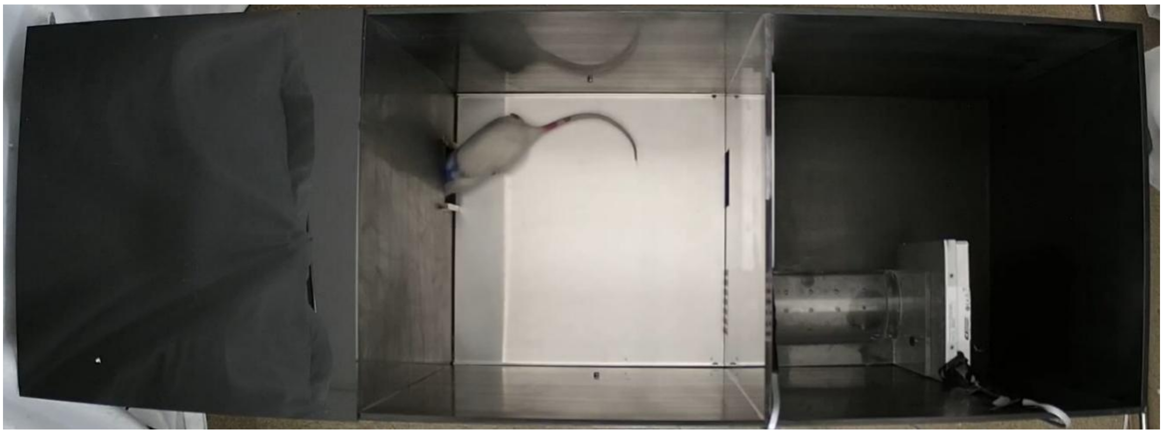

Figure S6. Experiment 2&3: the test phase for the free rat in the Non-social contact group (Empty restrainer condition)

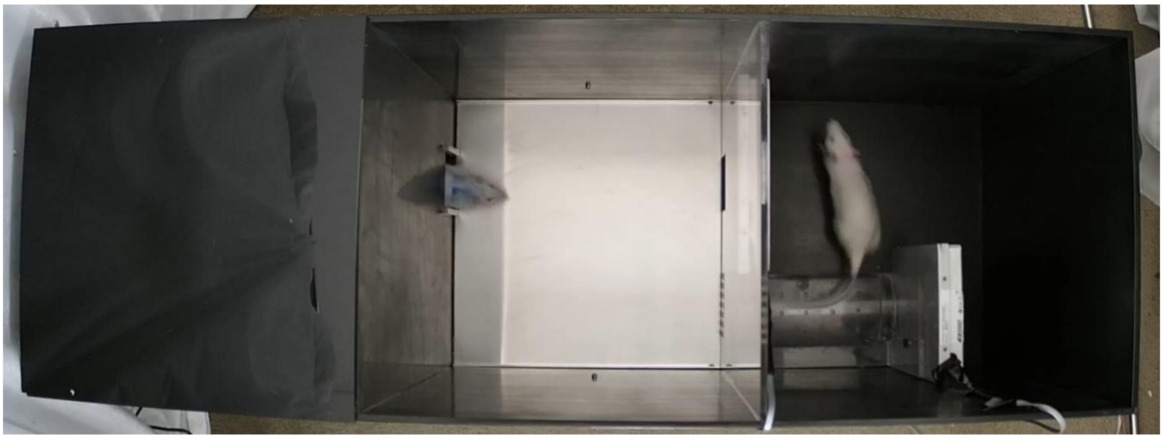

Figure S7. Experiment 2&3: the test phase for the free rat in the Non-social contact group (Familiar rat condition)

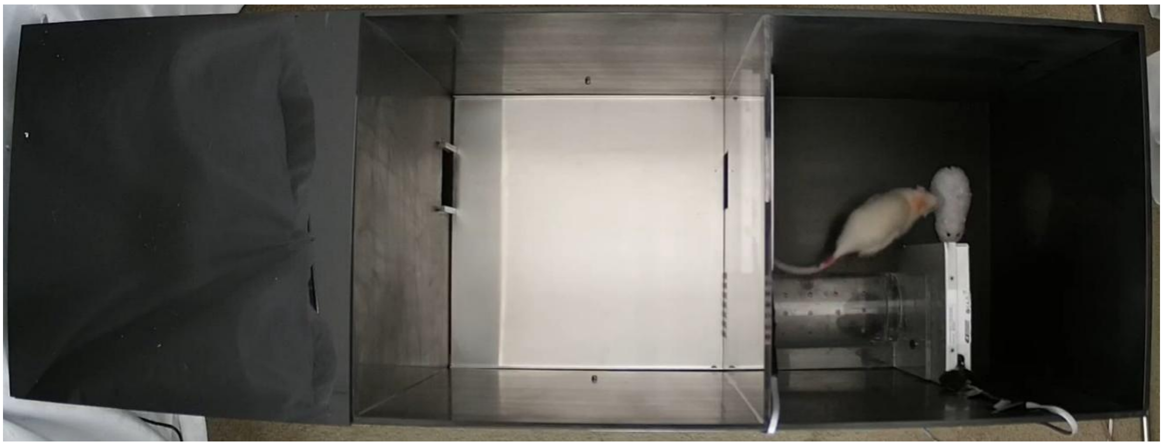

Figure S8. Experiment 2&3: the test phase for the free rat in the social contact group (Toy rat condition)

**Table S1.** Experiment 1: Inferential statistics of door-opening latency (s) in learning phase

| Factor | $F$ | $p$ | $\eta^2_{partial}$ |
|---|---|---|---|
| Experimental condition | 0.36 | .553 | .01 |
| Session | 2.96 | < .001 | .10 |
| Experimental condition × Session | 1.53 | .093 | .06 |

**Table S2-1.** Experiment 2: Inferential statistics of door-opening latency (s) in learning phase (Day 1 included)

| Factor | $F$ | $p$ | $\eta^2_{partial}$ |
|---|---|---|---|
| Experimental condition | 6.02 | .021 | .19 |
| Session | 9.40 | < .001 | .27 |
| Experimental condition × Session | 1.66 | .046 | .06 |

**Table S2-2.** Experiment 2: Inferential statistics of door-opening latency (s) in learning phase (Day 1 excluded)

| Factor | $F$ | $p$ | $\eta^2_{partial}$ |
|---|---|---|---|
| Experimental condition | 3.38 | .078 | .12 |
| Session | 9.50 | < .001 | .27 |
| Experimental condition × Session | 1.03 | .419 | .04 |

**Table S3.** Experiment 3: Inferential statistics of door-opening latency (s) in learning phase

| Factor | $F$ | $p$ | $\eta^2_{partial}$ |
|---|---|---|---|
| Social contact experience | 0.19 | .669 | .01 |
| Social contact in the experiment | 0.52 | .476 | .02 |
| Session | 4.74 | < .001 | .17 |
| Social contact experience × Social contact in the experiment | 1.38 | .251 | .05 |
| Session × Social contact experience | 3.50 | < .001 | .13 |
| Session × Social contact in the experiment | 0.56 | .905 | .02 |
| Session × Social contact experience × Social contact in the experiment | 0.71 | .772 | .03 |

**Table S4.** Experiment 1: Descriptive statistics of door-opening latency (s) in test phase

| **Experimental condition** | **Restrainer condition** | **Early period** | **Middle period** | **Late period** | **Average** | **Restrainer condition total average** |
|---|---|---|---|---|---|---|
| | | ***M±SE*** | ***M±SE*** | ***M±SE*** | ***M±SE*** | ***M±SE*** |
| Social contact group | Empty restrainer | 59.8±12.0 | 51.8±6.0 | 75.6±8.6 | 62.4±6.2 | 284.4±47.3 |
| | Familiar rat | 92.0±17.4 | 53.7±4.9 | 54.7±8.2 | 66.8±7.7 | 307.7±50.0 |
| | Unfamiliar rat | 61.1±8.5 | 54.0±6.3 | 54.1±4.0 | 56.4±4.6 | 303.6±52.1 |
| | Toy rat | 56.8±7.0 | 65.0±15.0 | 61.9±9.4 | 61.2±7.9 | 273.3±45.9 |
| | **Average** | 67.4±10.5 | 56.1±5.9 | 61.6±6.9 | | **61.7±5.9** |
| Non-social contact group | Empty restrainer | 269.9±38.8 | 516.5±62.8 | 732.6±48.9 | 506.3±40.9 | |
| | Familiar rat | 321.8±39.5 | 553.3±62.3 | 770.9±48.8 | 548.7±37.6 | |
| | Unfamiliar rat | 285.2±54.8 | 564.4±62.6 | 802.9±35.0 | 550.8±42.8 | |
| | Toy rat | 257.7±44.3 | 489.7±56.2 | 708.8±50.2 | 485.4±42.2 | |
| | **Average** | 283.6±39.3 | 531.0±55.8 | 753.8±36.7 | | **522.8±38.0** |
| **Period of test phase total average** | | 175.5±28.8 | 293.6±53.3 | 407.7±69.1 | | **292.3±48.2** |

**Table S5.** Experiment 1: Inferential statistics of door-opening latency (s) in test phase

| **Factor** | ***F*** | ***p*** | $\eta^2_{partial}$ |
|---|---|---|---|
| Experimental condition | 144.16 | < .001 | .85 |
| Period of test phase | 60.89 | < .001 | .70 |
| Condition of restrainer | 3.22 | .042 | .11 |
| Period of test phase × Experimental condition | 64.16 | < .001 | .71 |
| Experimental condition × Condition of restrainer | 3.27 | .040 | .11 |
| Period of test phase × Condition of restrainer | 0.60 | .689 | .02 |
| Period of test phase × Experimental condition × Condition of restrainer | 0.51 | .761 | .02 |

**Table S6.** Program Reversal Experiment： Descriptive statistics of door-opening latency (s) of social contact group and Non-social contact group under different conditions

| Session | Social contact group | | Non-social contact group | |
|---|---|---|---|---|
| | Empty restrainer | Familiar rat | Empty restrainer | Familiar rat |
| | *M*±*SE* | *M*±*SE* | *M*±*SE* | *M*±*SE* |
| 1 | 70.5±36.2 | 220.7±140.6 | 566.2±204.6 | 300.8±155.8 |
| 2 | 311.5±136.2 | 77.2±18.6 | 76.4±15.4 | 575.8±199.7 |
| 3 | 44.2±17.5 | 92.7±32.2 | 900.0±0 | 740.8±159.2 |
| 4 | 38.0±8.6 | 47.2±20.1 | 900.0±0 | 540.0±169.3 |
| 5 | 202.2±86.5 | 75.2±18.6 | 655.4±158.1 | 586.6±192.0 |
| 6 | 88.3±22.1 | 102.3±30.9 | 54.8±28.6 | 246.2±168.1 |
| 7 | 113.3±31.9 | 111.3±72.5 | 423.8±196.2 | 354.2±48.5 |
| 8 | 114.3±26.8 | 212.0±48.9 | 424.2±195.0 | 249.0±164.6 |
| 9 | 327.7±128.5 | 110.3±28.3 | 108.2±44.7 | 139.4±52.4 |
| 10 | 161.3±52.6 | 445.0±175.1 | 41.6±11.5 | 57.8±17.2 |
| 11 | 345.0±119.0 | 298.2±132.6 | 90.4±33.9 | 45.6±12.4 |
| 12 | 208.8±140.5 | 205.8±80.7 | 140.6±92.8 | 41.0±17.8 |
| 13 | 467.7±143.9 | 151.3±65.9 | 42.4±10.6 | 64.8±45.4 |
| 14 | 254.7±129.9 | 301.7±121.3 | 121.2±94.5 | 25.2±6.3 |
| 15 | 538.5±163.4 | 314.8±147.9 | 21.6±3.4 | 41.8±20.0 |

**Table S7.** Experiment 2: Descriptive statistics of door-opening latency (s) in test phase

| Experimental condition | Restrainer condition | Early period | Middle period | Late period | Average | Restrainer condition total average |
|---|---|---|---|---|---|---|
| | | *M*±*SE* | *M*±*SE* | *M*±*SE* | *M*±*SE* | *M*±*SE* |
| Social contact group | Empty restrainer | 82.7±11.7 | 76.9±15.5 | 85.0±10.1 | 81.5±11.4 | 221.4±31.0 |
| | Familiar rat | 120.1±16.4 | 69.4±8.1 | 71.2±9.6 | 86.9±8.4 | 280.4±42.7 |
| | Unfamiliar rat | 67.4±5.9 | 65.2±6.2 | 75.2±12.0 | 69.3±6.7 | 261.5±39.9 |
| | Toy rat | 57.3±5.1 | 66.6±7.5 | 81.7±15.4 | 68.5±8.1 | 201.1±31.4 |
| | **Average** | 81.9±7.4 | 69.5±7.9 | 78.3±11.2 | | **76.6±7.9** |
| Non-social contact group | Empty restrainer | 226.1±27.7 | 317.0±34.8 | 540.7±50.0 | 361.3±29.1 | |
| | Familiar rat | 308.6±43.6 | 389.4±61.4 | 724.0±41.7 | 474.0±41.5 | |
| | Unfamiliar rat | 213.7±31.4 | 458.8±46.3 | 688.8±36.6 | 453.8±29.6 | |
| | Toy rat | 195.3±34.0 | 272.1±37.8 | 533.9±56.9 | 333.8±36.5 | |
| | **Average** | 235.9±25.9 | 359.3±34.3 | 621.9±39.2 | | **405.7±28.5** |
| **Period of test phase total average** | | 158.9±19.9 | 214.4±32.8 | 350.1±56.0 | | **241.1±34.8** |

**Table S8.** Experiment 2: Inferential statistics of door-opening latency (s) in test phase

| **Factor** | ***F*** | ***p*** | $\eta^2_{partial}$ |
|---|---|---|---|
| Experimental condition | 123.75 | < .001 | .83 |
| Period of test phase | 77.10 | < .001 | .75 |
| Condition of restrainer | 9.97 | < .001 | .28 |
| Period of test phase × Experimental condition | 77.91 | < .001 | .75 |
| Experimental condition × Condition of restrainer | 8.17 | .001 | .24 |
| Period of test phase×Condition of restrainer | 3.61 | .006 | .12 |
| Period of test phase × Experimental condition × Condition of restrainer | 3.90 | .004 | .13 |

**Table S9.** Descriptive statistics of door-opening latency (s) of the social contact group between Experiment 1 and Experiment 2

| **Experiment** | **Period of test phase** | **Restrainer condition** | | | | **Experiment average** |
|---|---|---|---|---|---|---|
| | | **Empty restrainer** | **Familiar rat** | **Unfamiliar rat** | **Toy rat** | |
| | | ***M±SE*** | ***M±SE*** | ***M±SE*** | ***M±SE*** | ***M±SE*** |
| Experiment 1 | Early period | 59.8±12.0 | 92.0±17.4 | 61.1±8.5 | 56.8±7.0 | 61.7±5.9 |
| | Middle period | 51.8±6.0 | 53.7±4.9 | 54.0±6.3 | 65.0±15.0 | |
| | Late period | 75.6±8.6 | 54.7±8.2 | 54.1±4.0 | 61.9±9.4 | |
| Experiment 2 | Early period | 82.7±11.7 | 120.1±16.4 | 67.4±5.9 | 57.3±5.1 | 76.6±7.9 |
| | Middle period | 76.9±15.5 | 69.4±8.1 | 65.2±6.2 | 66.6±7.5 | |
| | Late period | 85.0±10.1 | 71.2±9.6 | 75.2±12.0 | 81.7±15.4 | |

**Table S10.** Inferential statistics of door-opening latency (s) of the social contact group between Experiment 1 and Experiment 2

| **Factor** | ***F*** | ***p*** | $\eta^2_{partial}$ |
|---|---|---|---|
| Experiment | 2.27 | .144 | .08 |
| Period of test phase | 1.95 | .152 | .07 |
| Condition of restrainer | 4.87 | .004 | .16 |
| Period of test phase×Experiment | 0.04 | .961 | .00 |
| Experiment×Condition of restrainer | 1.04 | .380 | .04 |
| Period of test phase×Condition of restrainer | 9.40 | < .001 | .27 |
| Period of test phase×Experiment×Condition of restrainer | 1.06 | .386 | .04 |

**Table S11.** Descriptive statistics of door-opening latency (s) of the Non-social contact group between Experiment 1 and Experiment 2

| Experiment | Period of test phase | Restrainer condition | | | | Experiment average |
|---|---|---|---|---|---|---|
| | | Empty restrainer | Familiar rat | Unfamiliar rat | Toy rat | |
| | | *M±SE* | *M±SE* | *M±SE* | *M±SE* | *M±SE* |
| Experiment 1 | Early period | 269.9±38.8 | 321.8±39.5 | 285.2±54.8 | 257.7±44.3 | 522.8±38.0 |
| | Middle period | 516.5±62.8 | 553.3±62.3 | 564.4±62.6 | 489.7±56.2 | |
| | Late period | 732.6±48.9 | 770.9±48.8 | 802.9±35.0 | 708.8±50.2 | |
| Experiment 2 | Early period | 226.1±27.7 | 308.6±43.6 | 213.7±31.4 | 195.3±34.0 | 405.7±28.5 |
| | Middle period | 317.0±34.8 | 389.4±61.4 | 458.8±46.3 | 272.1±37.8 | |
| | Late period | 540.7±50.0 | 724.0±41.7 | 688.8±36.6 | 533.9±56.9 | |

**Table S12.** Inferential statistics of door-opening latency (s) of the Non-social contact group between Experiment 1 and Experiment 2

| Factor | $F$ | $p$ | $\eta^2_{partial}$ |
|---|---|---|---|
| Experiment | 6.09 | .021 | .19 |
| Period of test phase | 140.21 | < .001 | .84 |
| Condition of restrainer | 12.31 | < .001 | .32 |
| Period of test phase × Experiment | 3.05 | .056 | .11 |
| Experiment × Condition of restrainer | 1.70 | .175 | .06 |
| Period of test phase × Condition of restrainer | 2.62 | .019 | .09 |
| Period of test phase × Experiment × Condition of restrainer | 1.12 | .351 | .04 |

**Table S13-1.** Experiment 3: Descriptive statistics of door-opening latency (s) in test phase (Detailed version)

| Experimental condition | | Period of test phase | Empty restrainer | Familiar rat | Unfamiliar rat | Toy rat | Average | Experimental condition total average |
|---|---|---|---|---|---|---|---|---|
| | | | ***M*±*SE*** | ***M*±*SE*** | ***M*±*SE*** | ***M*±*SE*** | ***M*±*SE*** | ***M*±*SE*** |
| Social contact condition | Social contact experience | Early period | 157.6±51.6 | 191.3±45.3 | 141.6±39.2 | 105.4±26.2 | 149.0±38.5 | |
| | | Middle period | 125.8±19.9 | 160.8±52.3 | 106.1±21.4 | 125.2±27.4 | 129.5±28.3 | |
| | | Late period | 115.1±21.0 | 85.3±14.8 | 94.3±12.6 | 134.1±37.9 | 107.2±12.3 | |
| | | **Average** | 132.8±21.9 | 145.8±32.5 | 114.0±22.9 | 121.6±20.6 | | **128.5±22.6** |
| | Non-social contact experience | Early period | 107.6±20.5 | 148.6±88.2 | 94.4±15.2 | 83.9±14.1 | 108.6±18.2 | |
| | | Middle period | 81.4±15.8 | 64.4±12.0 | 79.8±13.5 | 111.1±37.1 | 84.1±14.4 | |
| | | Late period | 62.9±5.4 | 98.4±19.2 | 59.5±11.1 | 80.0±16.7 | 75.2±10.7 | |
| | | **Average** | 84.0±9.2 | 103.8±15.0 | 77.9±10.3 | 91.6±12.5 | | **89.3±9.6** |
| | **Sub-total average** | | 108.4±13.3 | 124.8±18.2 | 95.9±13.1 | 106.6±12.3 | | **108.9±13.0** |
| Non-social contact condition | Social contact experience | Early period | 198.5±53.8 | 217.5±41.5 | 137.1±32.2 | 176.9±55.3 | 182.5±27.8 | |
| | | Middle period | 347.2±99.3 | 332.9±82.3 | 289.2±64.3 | 325.7±67.2 | 323.8±71.1 | |
| | | Late period | 486.4±99.6 | 444.4±56.9 | 509.1±86.5 | 467.4±100.7 | 476.8±53.2 | |
| | | **Average** | 344.1±60.7 | 331.6±43.1 | 311.8±49.5 | 323.4±47.1 | | **327.7±42.4** |
| | Non-social contact experience | Early period | 315.3±99.4 | 270.2±258.1 | 247.6±85.4 | 309.6±104.2 | 285.7±91.0 | |
| | | Middle period | 516.6±118.7 | 533.4±130.4 | 558.9±111.8 | 555.1±129.8 | 541.0±118.7 | |
| | | Late period | 459.6±110.6 | 550.8±105.0 | 557.5±132.6 | 583.7±124.7 | 537.9±100.6 | |
| | | **Average** | 430.5±102.2 | 451.5±100.0 | 454.7±96.4 | 482.8±95.7 | | **454.9±93.7** |
| | **Sub-total average** | | 387.3±58.4 | 391.5±54.9 | 383.2±55.7 | 403.1±55.8 | | **391.3±52.5** |
| **Restrainer condition total average** | | | 247.8±39.8 | 258.2±38.3 | 239.6±39.4 | 254.8±40.0 | | **250.1±38.0** |

**Table S13-2.** Experiment 3: Descriptive statistics of door-opening latency (s) in test phase (Aggregated version)

| Experimental condition | Early period | Middle period | Late period | Experimental condition average (*M*±*SE*) |
|---|---|---|---|---|
| | *M*±*SE* | *M*±*SE* | *M*±*SE* | |
| Social contact condition | 128.8±21.2 | 106.8±16.5 | 91.2±9.0 | 108.9±13.0 |
| Non-social contact condition | 234.1±47.9 | 432.4±73.0 | 507.4±55.3 | 391.3±52.5 |
| Social contact experience | 165.7±23.3 | 226.6±45.6 | 292.0±57.6 | 228.1±36.0 |
| Non-social contact experience | 197.2±50.9 | 312.6±85.5 | 306.6±80.5 | 272.1±68.0 |
| Period of test phase average | 181.4±27.6 | 269.6±48.3 | 299.3±48.6 | |

**Table S14.** Experiment 3: Inferential statistics of door-opening latency (s) in test phase

| Factor | *F* | *p* | $\eta^2_{partial}$ |
|---|---|---|---|
| Experimental condition | 28.50 | < .001 | .54 |
| Social contact experience | 0.69 | .414 | .03 |
| Period of test phase | 11.24 | < .001 | .32 |
| Restrainer condition | 0.44 | .727 | .02 |
| Social contact experience × Experimental condition | 2.47 | .129 | .09 |
| Period of test phase × Experimental condition | 19.12 | < .001 | .44 |
| Period of test phase × Social contact experience | 1.04 | .361 | .04 |
| Experimental condition × Restrainer condition | 0.26 | .855 | .01 |
| Period of test phase × Restrainer condition | 0.83 | .549 | .03 |
| Social contact experience × Restrainer condition | 0.64 | .595 | .03 |
| Period of test phase × Experimental condition × Restrainer condition | 0.59 | .740 | .03 |
| Period of test phase × Social contact experience × Experimental condition | 1.42 | .252 | .06 |
| Restrainer condition × Social contact experience × Experimental condition | 0.23 | .878 | .01 |
| Restrainer condition × Social contact experience × Period of test phase | 0.68 | .663 | .03 |
| Restrainer condition × Social contact experience × Period of test phase × Experimental condition | 0.36 | .901 | .02 |

**Table S15.** Descriptive statistics of door-opening latency (s) of the social contact group between Experiment 2 and Experiment 3

| Experiment | Period of test phase | Restrainer condition | | | | Experiment total |
|---|---|---|---|---|---|---|
| | | Empty restrainer | Familiar rat | Unfamiliar rat | Toy rat | |
| | | *M*±*SE* | *M*±*SE* | *M*±*SE* | *M*±*SE* | *M*±*SE* |
| Experiment 2 | Early period | 86.2±16.0 | 148.0±23.1 | 70.2±6.1 | 59.8±5.8 | 75.7±7.0 |
| | Middle period | 78.4±15.2 | 68.5±8.0 | 64.8±7.1 | 55.0±5.8 | |
| | Late period | 74.7±14.1 | 67.6±9.0 | 64.0±7.1 | 71.0±9.1 | |
| Experiment 3 | Early period | 132.6±27.6 | 169.9 ±27.6 | 118.0±21.2 | 94.6±14.6 | 108.9±13.0 |
| | Middle period | 103.6±13.7 | 112.6±29.0 | 93.0±12.7 | 118.1±22.2 | |
| | Late period | 89.0±12.7 | 91.9±11.8 | 76.9±9.4 | 107.0±21.2 | |

**Table S16.** Inferential statistics of door-opening latency (s) of the social contact group between Experiment 2 and Experiment 3

| Factor | *F* | *p* | $\eta^2_{partial}$ |
|---|---|---|---|
| Experiment | 5.08 | .033 | .16 |
| Period of test phase | 5.53 | .007 | .18 |
| Condition of restrainer | 7.37 | < .001 | .22 |
| Period of test phase × Experiment | 0.56 | .574 | .02 |
| Experiment × Condition of restrainer | 0.65 | .586 | .02 |
| Period of test phase × Condition of restrainer | 6.51 | < .001 | .20 |
| Period of test phase × Experiment × Condition of restrainer | 0.76 | .606 | .03 |

**Table S17.** Descriptive statistics of door-opening latency (s) of the Non-social contact group between Experiment 2 and Experiment 3

| Experiment | Period of test phase | Restrainer condition | | | | Experiment total |
|---|---|---|---|---|---|---|
| | | Empty restrainer | Familiar rat | Unfamiliar rat | Toy rat | |
| | | *M*±*SE* | *M*±*SE* | *M*±*SE* | *M*±*SE* | *M*±*SE* |
| Experiment 2 | Early period | 182.4±25.0 | 285.5±46.6 | 178.6±23.9 | 169.1±30.5 | 297.6±28.1 |
| | Middle period | 302.6±35.0 | 363.5±57.8 | 350.9±50.4 | 195.0±30.5 | |
| | Late period | 329.6±42.4 | 398.0±58.9 | 479.3±46.6 | 336.9±49.3 | |
| Experiment 3 | Early period | 256.9±56.7 | 243.9±51.4 | 192.4±46.4 | 243.3±59.6 | 391.3±52.5 |
| | Middle period | 431.9±78.0 | 433.1±79.1 | 424.1±72.4 | 440.4±77.0 | |
| | Late period | 473.0±71.6 | 497.6±59.2 | 533.3±76.3 | 525.6±78.7 | |

**Table S18.** Inferential statistics of door-opening latency (s) of the Non-social contact group between Experiment 2 and Experiment 3

| Factor | $F$ | $p$ | $\eta^2_{partial}$ |
|---|---|---|---|
| Experiment | 2.48 | .128 | .09 |
| Period of test phase | 36.50 | < .001 | .58 |
| Condition of restrainer | 2.13 | .103 | .08 |
| Period of test phase × Experiment | 2.07 | .136 | .07 |
| Experiment × Condition of restrainer | 3.25 | .026 | .11 |
| Period of test phase × Condition of restrainer | 2.26 | .040 | .08 |
| Period of test phase × Experiment × Condition of restrainer | 0.48 | .823 | .02 |

**Table S19.** Descriptive statistics of duration (s) in the dark chamber under different Experimental condition

| Experimental condition | Place | Empty restrainer | Familiar rat | Unfamiliar rat | Toy rat | Total average |
|---|---|---|---|---|---|---|
| | | ***M±SE*** | ***M±SE*** | ***M±SE*** | ***M±SE*** | ***M ± SE*** |
| Social contact | Dark Chamber | 25.8±4.5 | 24.0±3.5 | 17.6±2.4 | 20.3±3.2 | 21.9±2.9 |
| | Middle Chamber | 55.7±7.2 | 62.8±5.1 | 51.7±5.5 | 48.2±5.4 | 54.6±5.2 |
| Non-social contact | Dark Chamber | 81.0±9.8 | 116.4±12.3 | 102.2±8.2 | 56.1±4.9 | 88.9±5.5 |
| | Middle Chamber | 280.2±29.2 | 357.6±37.0 | 351.6±27.4 | 277.7±33.5 | 316.8±27.4 |
